# Disentangling the neural correlates of semantic and domain-general control: The roles of stimulus domain and task process

**DOI:** 10.1101/2023.08.23.554418

**Authors:** Victoria J. Hodgson, Matthew A. Lambon Ralph, Rebecca L. Jackson

**Affiliations:** MRC Cognition and Brain Sciences Unit, University of Cambridge, UK; Department of Psychology & York Biomedical Research Institute, University of York, UK

**Keywords:** Control, Semantic control, Multiple demand network, Language, Executive function, Working memory

## Abstract

Control processes are critical for the context-appropriate use of meaningful stimuli. Similar definitions have been adopted in two distinct literatures focusing on identifying the neural correlates of ‘semantic control’ and of executive control across domains (the ‘multiple demand network’). Surprisingly, despite their proposed functions varying only in relation to domain-specificity, these networks appear to differ anatomically. However, prior comparisons are confounded by variations in task design. To what extent might varying task requirements drive differences in activation patterns that are typically attributed to stimulus domain? Here, for the first time, we use functional MRI to disentangle the effects of task process and stimulus domain during cognitively demanding tasks. Participants performed an odd-one-out task requiring rule-switching, inhibition and selection processes, and an *n*-back working memory task, each with meaningful semantic and non-semantic stimuli, in a factorial design. Both stimulus domain and task process affected the control regions activated, indicating that task process is indeed a key factor confounding prior studies. However, core semantic control regions (left inferior frontal gyrus, left posterior temporal cortex) also showed a preference for semantic stimuli even with matched task processes, while more peripheral semantic control regions, overlapping the multiple demand network (dorsomedial prefrontal cortex, right inferior frontal gyrus), showed little preference across task or stimulus. Conversely, most multiple demand network regions were preferentially engaged for non-semantic stimuli. These results highlight the mutual importance of stimulus domain and task process in driving variation in control region engagement, both across and between semantic control and multiple demand networks.

**SIGNIFICANCE STATEMENT:** The flexible, context-appropriate use of concepts requires the selection, inhibition and manipulation of meaningful information. These control processes are thought to be supported by different areas for conceptual processing compared to other task domains. This proposed ‘special’ character of semantic control has important ramifications for the nature of executive control. However, prior assessments confound the presence of meaningful stimuli with the task operations performed. Here we disentangle the effects of task process and stimulus domain for the first time, finding critical effects of both factors on the pattern of activated control regions. The results enhance our understanding of the semantic control network and how it differs from and interacts with the domain-general multiple demand network, functionally characterising each control region.

## INTRODUCTION

Semantic knowledge is represented through the interaction of a multimodal hub in the bilateral anterior temporal lobes (ATL) and distributed sensory-specific spoke regions ^1–3^. However, representation processes alone are insufficient for successful semantic cognition. Flexible, semantically-driven behaviour requires semantic control: the effortful, context-dependent manipulation and selection of meaningful semantic information for the purposes of completing a task or goal ^1,4^. Semantic control demands are high in tasks that require selecting non-dominant features, identifying weak semantic associations, inhibiting task-irrelevant features, resolving ambiguity or task-switching ^1,5–10^. This control of meaningful information is subserved by a left-lateralised network of inferior frontal and posterior temporal regions distinct from semantic representation areas, known as the semantic control network (SCN). Evidence for the critical nature of semantic control initially came from patients with semantic aphasia, who display deficits in the control of conceptual information following a stroke affecting left prefrontal or temporoparietal cortex ^11–13^. Crucially, these deficits are dependent on task context ^11,12^ and qualitatively distinct from those observed in semantic dementia, in which degraded semantic representations from bilateral ATL atrophy cause consistent deficits for particular concepts across contexts ^5,12–19^. Subsequent neuroimaging and transcranial magnetic stimulation studies confirmed the involvement of inferior frontal gyrus (IFG; particularly on the left) and left posterior temporal cortex in controlled, semantic processing, with neuroimaging additionally highlighting a role for bilateral dorsomedial prefrontal cortex (dmPFC) in and around the presupplementary motor area ^4,5,7,9,20–23^.

The SCN is typically only studied in the context of the semantic literature, yet numerous domains require control processes. Indeed, the multiple demand network (MDN) is considered to underpin executive control regardless of domain, having been defined based on the convergence of fMRI activation across many different effortful cognitive tasks ^24,25^. A similar set of cognitive processes are proposed to be supported by the MDN and the SCN, including selective attention and retrieval of goal-relevant knowledge, inhibition of dominant yet task-irrelevant information and deciding between competing alternatives ^1,4,21,24,26–29^. The crucial difference in the definitions of the two networks is their proposed domain-specificity. Thus, by considering the relationship between measures of the MDN and SCN, we may ask whether semantic tasks simply recruit domain-general control areas or whether the SCN reflects an additional, distinct resource. Note that the two networks are defined independently, meaning areas involved in multiple domains are not excluded from the SCN, therefore areas involved in the executive control of all domains including semantic cognition should be identified in both.

The MDN comprises bilateral frontal and parietal regions, including dmPFC, premotor cortex, middle frontal gyrus (MFG), anterior insula, and inferior parietal lobe (IPL) ^24,26,30–32^. In recent investigations, these core areas have been expanded to include ‘extended’ MDN regions in posterior inferior temporal cortex, frontal pole and premotor cortex ^31,33^. Thus, while there are regions that may be identified in both the MDN and SCN including dmPFC, left IFG and left posterior inferior temporal gyrus (pITG) ^4,24,26,34^, these are within the context of substantial differences in the extent and foci of the two networks ^4,35–37^. In lateral frontal cortex, core MDN areas are more dorsal (including superior frontal gyrus (SFG), MFG and dorsal IFG) than the IFG-focused SCN ^4,20,24,26,33,38,39^. Parietal cortex is not reliably activated across studies of semantic control ^4^, yet IPL is a core region of the MDN ^26,33^. Left posterior temporal cortex also appears to show heterogeneity in its domain-specificity; while the extended MDN includes some pITG, the SCN includes a broad swathe of activation across posterior superior temporal sulcus (pSTS), posterior middle temporal gyrus (pMTG) and pITG ^4,9^.

Given the apparent similarity of the processes required for the control of semantic and non-semantic domains, why, then, do the SCN and MDN not overlap more closely? One possibility is that meaning may have some ‘special’ status whereby its presence necessitates the recruitment of specialised neural control mechanisms (either instead of, or in addition to, domain-general control regions). However, variation in domain-specificity may not be the only explanation for these observed differences. Methods used to define popular templates of each network differ, with the SCN delineated using meta-analyses ^4,9^ and the MDN requiring overlapping activity across multiple tasks ^26,40^. Furthermore, both these broad cross-study assessments and direct comparisons tend to conflate the presence of meaningful stimuli with the type of cognitive process required. For instance, studies contrasting semantic and domain-general control have compared meaningful stimuli using word association judgements or feature-matching to meaningless stimuli utilising working memory, visuospatial reasoning or inhibition and attention tasks ^20,35–37,41–43^. Similarly, the SCN meta-analyses may have a greater focus on particular task processes, such as selection and ambiguity resolution, and less involvement of others, such as working memory, task-switching or attention, than typical MDN-focused experiments. Therefore, it is difficult to dissociate the effects of stimulus domain and task process within the existing body of literature.

In the present fMRI experiment, we delineate the independent effects of task process and stimulus domain, as well as their interaction, on the regions recruited for executive control. We employ a factorial design with two distinct, yet commonly utilised, tasks designed to tap into different control processes, and two sets of stimuli to assess semantic and non-semantic domains within a single participant sample. The first task, an extension of the Adapted Cattell Culture Fair task from Woolgar et al. ^28^ (itself based on the Cattell Culture Fair test battery ^44^), requires participants to identify the odd-one-out from an array of items, by selecting and inhibiting different stimulus features to generate different potential rules and switch flexibly between these candidate rules. These processes are considered central to the function of the MDN and, as such, this task (using non-semantic stimuli) has been used to delineate the MDN and demonstrate its relation to fluid intelligence ^27,28^. These processes are also considered core elements of semantic control and this type of semantic judgement is typical of semantic control studies. Therefore, a new variant utilising semantic stimuli was created here. As participants must perform the same kinds of executive processes to solve both variants, this allows comparison of the control regions activated across semantic and non-semantic stimulus domains without a confound of task process. The addition of a second task paradigm, the *n*-back working memory task with variants using semantic and non-semantic stimuli, also allows comparison between task processes without varying stimulus domain. Crucially, this paradigm targets working memory and attention, different aspects of executive function to the odd-one-out task ^40,45–49^, and activates MDN regions ^27,33,40,45,50–52^. This study design (shown in Figure 1) therefore allowed us to determine the independent effects of task process and stimulus domain on the pattern of regions recruited for control processing (which were compared to previous measures of the SCN and MDN) and the functional profile of individual control regions.

**Figure 1.**
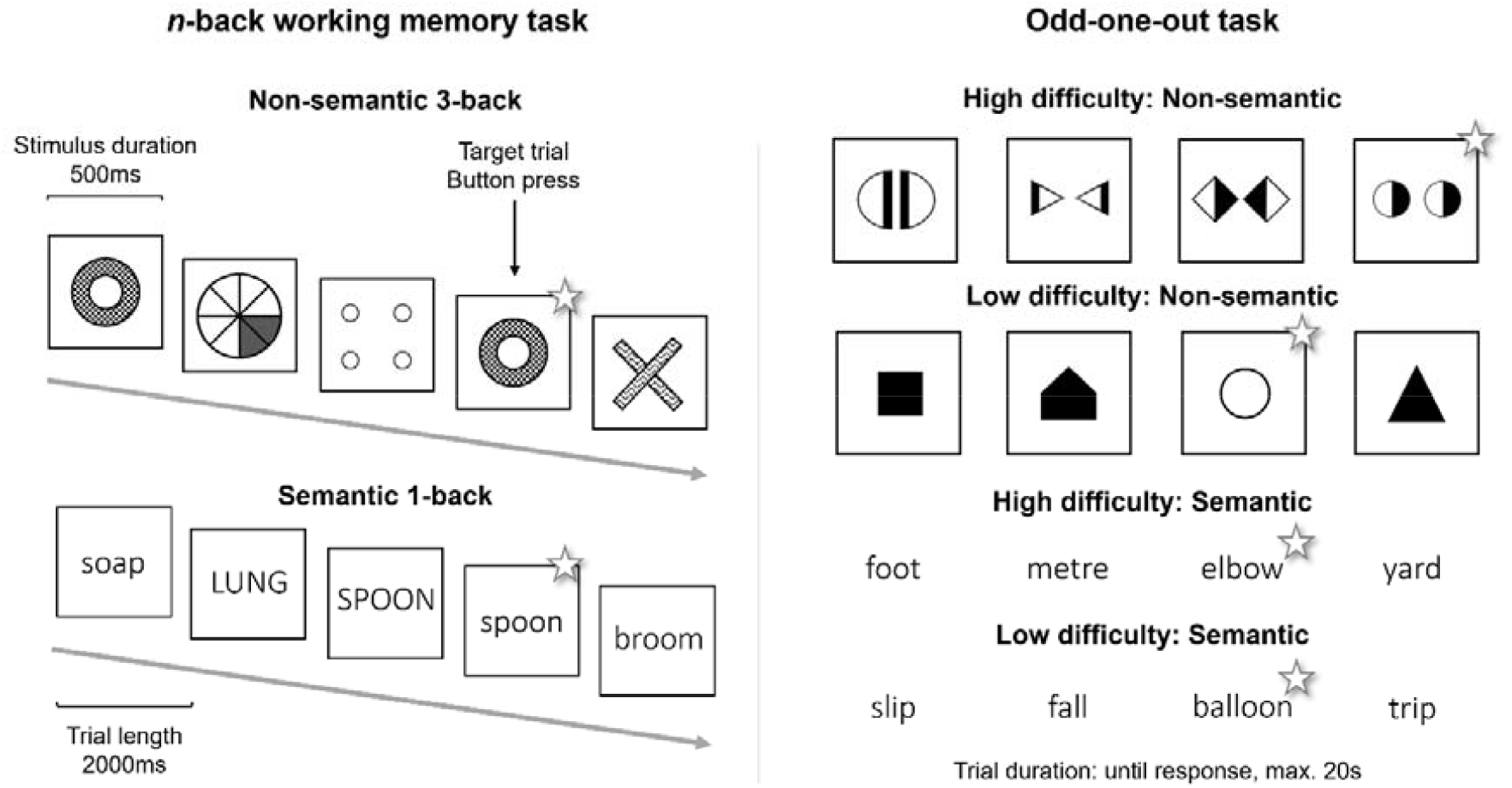
Illustration of the task design including the two stimulus types (semantic and non-semantic) and the two tasks (the odd-one-out and the n-back). Correct answers are indicated with a star. Left: the 3-back task with non-semantic stimuli (above) and the 1-back task with semantic stimuli (below). Each participant also performed a 3-back task with semantic stimuli and a 1-back task with non-semantic stimuli (not pictured). Right: all four variants of the odd-one-out task, adapted from the Cattell Culture Fair test battery ^44^ and Woolgar et al ^28^. Note, throughout the conditions are labelled ‘semantic’ or ‘non-semantic’ n-back or odd-one-out task. This refers to the presence of semantic or non-semantic stimuli only and not the processes used to perform the task. For instance, the ‘semantic n-back’ does not necessarily require in-depth semantic processing and could be solved using phonological working memory. However, an executive control process is applied to meaningful stimuli.

## RESULTS

### Behavioural data

Behavioural performance is shown in Table 1 (also see Supplementary Figure 1). Paired t-tests indicated no significant difference in accuracy or reaction time between the different hard or different easy variants of the odd-one-out task (all p values<0.05; see Supplementary Table 1), except the reaction time of the easy semantic and non-semantic variants which differed significantly, t(31) = 6.53, p<0.001. Thus, the easy semantic and non-semantic variants of the odd-one-out task were not well matched and comparing the hard>easy conditions of each would result in a much greater reaction time difference for the semantic than non-semantic variant, t(31) = -4.96, p<0.001. Therefore, difficulty effects would confound the comparison between semantic and non-semantic control when focusing on the hard>easy (but not hard>rest) contrast.

**Table 1.** Summary of behavioural data (32 participants)

| Difficulty | Condition | % Accuracy | d-prime | Reaction time (s) |
| --- | --- | --- | --- | --- |
| Hard | Semantic odd-one-out | 74.0 ± 9.54 | - | 5.84 ± 1.01 |
| | Non-semantic odd-one-out | 74.5 ± 8.56 | - | 5.98 ± 1.29 |
| | Semantic $n$-back | - | 2.81 ± 0.679 | 0.979 ± 0.0403 |
| | Non-semantic $n$-back | - | 2.05 ± 0.686 | 0.968 ± 0.0432 |
| Easy | Semantic odd-one-out | 95.1 ± 3.10 | - | 3.32 ± 0.960 |
| | Non-semantic odd-one-out | 92.6 ± 8.53 | - | 2.49 ± 0.592 |
| | Semantic $n$-back | - | 4.02 ± 0.700 | 0.676 ± 0.131 |
| | Non-semantic $n$-back | - | 3.72 ± 0.780 | 0.651 ± 0.125 |

Performance in the *n*-back task was assessed using d-prime ^53^. Paired t-tests indicated significant differences in d-prime between the semantic and non-semantic variants at both the hard (t(31) = -6.15, p<0.001) and easy levels (t(31) = -2.68, p=0.0118), though their reaction time did not differ. The non-semantic variant of the *n*-back task was somewhat more challenging than the semantic variant, for both hard and easy conditions, despite attempts to match these during behavioural piloting. Thus, some difficulty effects could be present in the semantic vs. non-semantic contrast within the *n*-back task.

### Whole-brain analyses

For each of the four task-stimulus combinations, the hard and easy conditions are each displayed over rest, alongside the difference between the hard and easy conditions (see Figure 2, Supplementary Table 2 for peaks). For both variants of the odd-one-out task, both easy>rest and hard>rest contrasts reveal a similar pattern of activation across frontal, parietal and temporal control regions. The strong engagement of control networks in the easy condition raises the possibility that activation in control networks is obscured in the hard>easy contrasts. Furthermore, the behavioural data indicated that the semantic and non-semantic easy conditions were not well-matched, therefore the extent to which this control network activation is subtracted out through comparison with the easy baseline is likely to differ across conditions. All further analyses therefore focus on the hard>rest data to promote the identification of the control areas and remove the confounding effect of difficulty between the semantic and non-semantic odd-one-out variants. However, the hard>easy contrasts in Figure 2 give a similar pattern of results (confirming the control-related interpretation of the regions identified, consistent with their identification in prior measures of the MDN and SCN).

**Figure 2.**
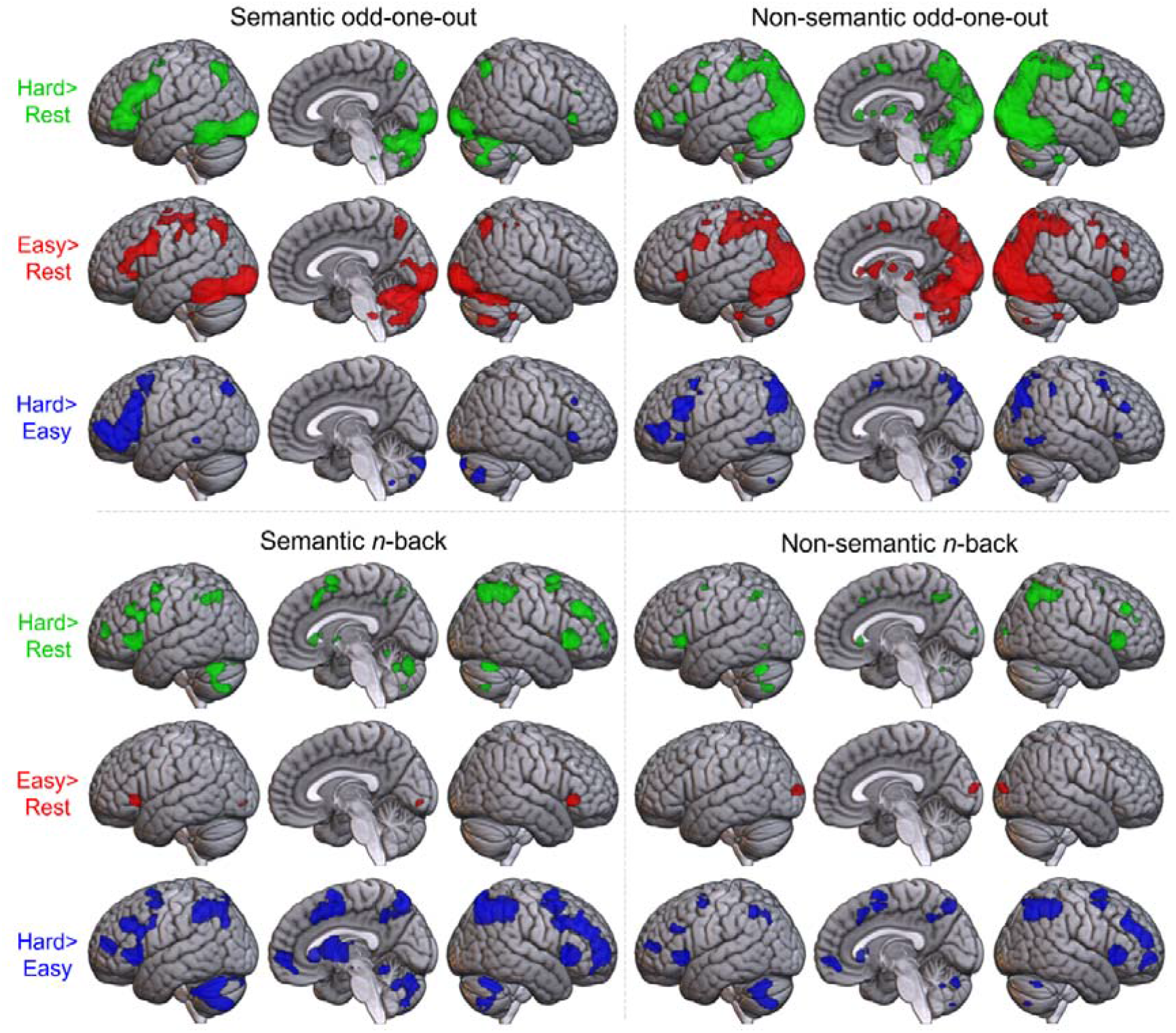
Results of univariate contrasts across all task-stimulus combinations. Each condition is shown in a different quadrant. Hard>easy (blue), hard>rest (green) and easy>rest (red) contrasts are shown for each task-stimulus condition. Voxel-level threshold p<0.05 with FWE correction.

Comparing the hard condition with rest highlighted a broadly similar pattern across the four conditions, with activation across lateral frontal cortex and the IPL. Importantly, in addition, there were clear differences between the four task-stimulus pairings. Both variants of the odd-one-out task had additional activation in posterior inferior temporal and occipital lobes. In the non-semantic odd-one-out task, a single large cluster spanned a large portion of bilateral occipital lobe, extending anteriorly into ventral posterior temporal lobe (inferior temporal and fusiform gyri) and the IPL. Only the semantic odd-one-out task activity appeared strongly left-lateralised, with large clusters in left IFG and left inferior temporal cortex – notably, key regions of the SCN – and less or no activation in their right hemisphere homologues. The semantic *n*-back task showed multiple, small clusters across the IPL, IFG and MFG bilaterally, as well as a medial cluster in dmPFC, including the supplementary motor area (SMA) and pre-SMA. The non-semantic *n*-back task revealed clusters in similar locations to the semantic *n*-back, but with a reduced extent.

The activation pattern in some conditions appears to better resemble the MDN or SCN. To quantify this, the overlap between the activity pattern in each condition and existing masks of the SCN (from Jackson ^4^) and MDN (from Fedorenko et al., ^26^) was assessed (Figure 3). The semantic odd-one-out task best reflected the SCN, while the non-semantic odd-one-out task strongly resembled the MDN. This demonstrates i) the strong alignment between the patterns of activity identified here and those present in the existing literature, and ii) that both task process and stimuli type can affect the control network identified. The alignment of the SCN with the semantic odd-one-out task is highly consistent with its proposed role in the controlled selection and manipulation of semantic stimuli. The MDN also appears best identified with a task focusing on these same processes, yet with non-semantic stimuli. While both *n*-back conditions include multiple regions associated with the MDN, this overlap is less extensive than the non-semantic odd-one-out condition due to the reduced involvement of the bilateral posterior occipital, parietal and temporal regions.

**Figure 3.**
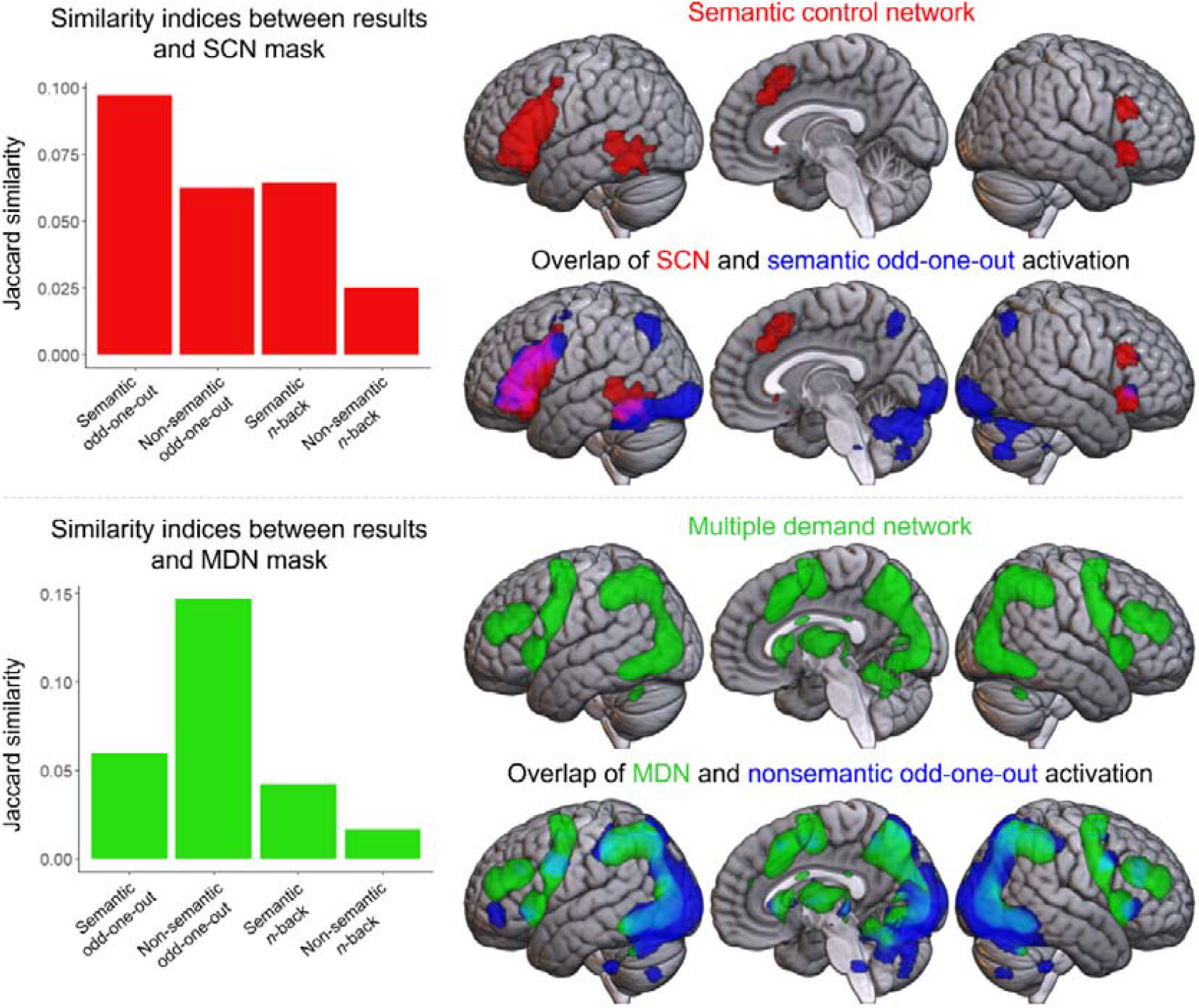
Results of the network overlap analysis. Left column: graphs showing the overlap between the pattern of activity in each condition and existing definitions of the SCN (top) and MDN (bottom) as measured using the Jaccard similarity index. For the SCN mask, the highest similarity index is seen for the semantic odd-one-out condition; for the MDN mask, the highest similarity index is seen for the non-semantic odd-one-out condition. Note, the reduced overlap with the MDN for the non-semantic than semantic n-back condition is due to a decreased extent of activation not a different activation profile and should be interpreted cautiously. Right column: each network template mask is shown independently (red for the SCN, green for the MDN) and overlapping with the condition with the highest Jaccard similarity index (indicated in each case in blue). Regions of overlap are violet (for SCN and semantic odd-one-out) or cyan (for MDN and non-semantic odd-one-out).

To explore the differences between conditions formally, a factorial two-way ANOVA was performed on the hard>rest data with the factors task process (odd-one-out vs. *n*-back) and stimulus domain (semantic vs. non-semantic). Results are displayed in Figure 4 (with peaks in Supplementary Table 3). There were extensive, bilateral main effects of task process, involving the majority of bilateral occipital lobe, pITG, IPL and anterior cingulate cortex, right STS, left IFG and a smaller region of right IFG and insula. Post hoc comparisons showed greater activation for the odd-one-out than *n-*back task in left IFG (pars triangularis and orbitalis), right IFG (pars triangularis), and a large bilateral swathe encompassing inferior occipital cortex and extending anteriorly into pITG/fusiform gyrus, and dorsally into the parietal lobe and PCG. The *n*-back task showed greater activation in bilateral MFG, insula, precuneus and posterior and anterior cingulate, right IFG and left AG and pSTG. There was also reduced deactivation for the *n*-back task in right temporo-parietal junction, including right AG and right mid and posterior STS. There are broad effects of task throughout the control networks, including key areas thought to differ between semantic and domain-general control, such as MFG, IFG and left pITG. Thus, task effects are likely to have contributed to the differences identified in prior comparisons of semantic and domain-general control. However, this is not the only factor underpinning these differences; there are also effects of stimulus domain.

**Figure 4.**
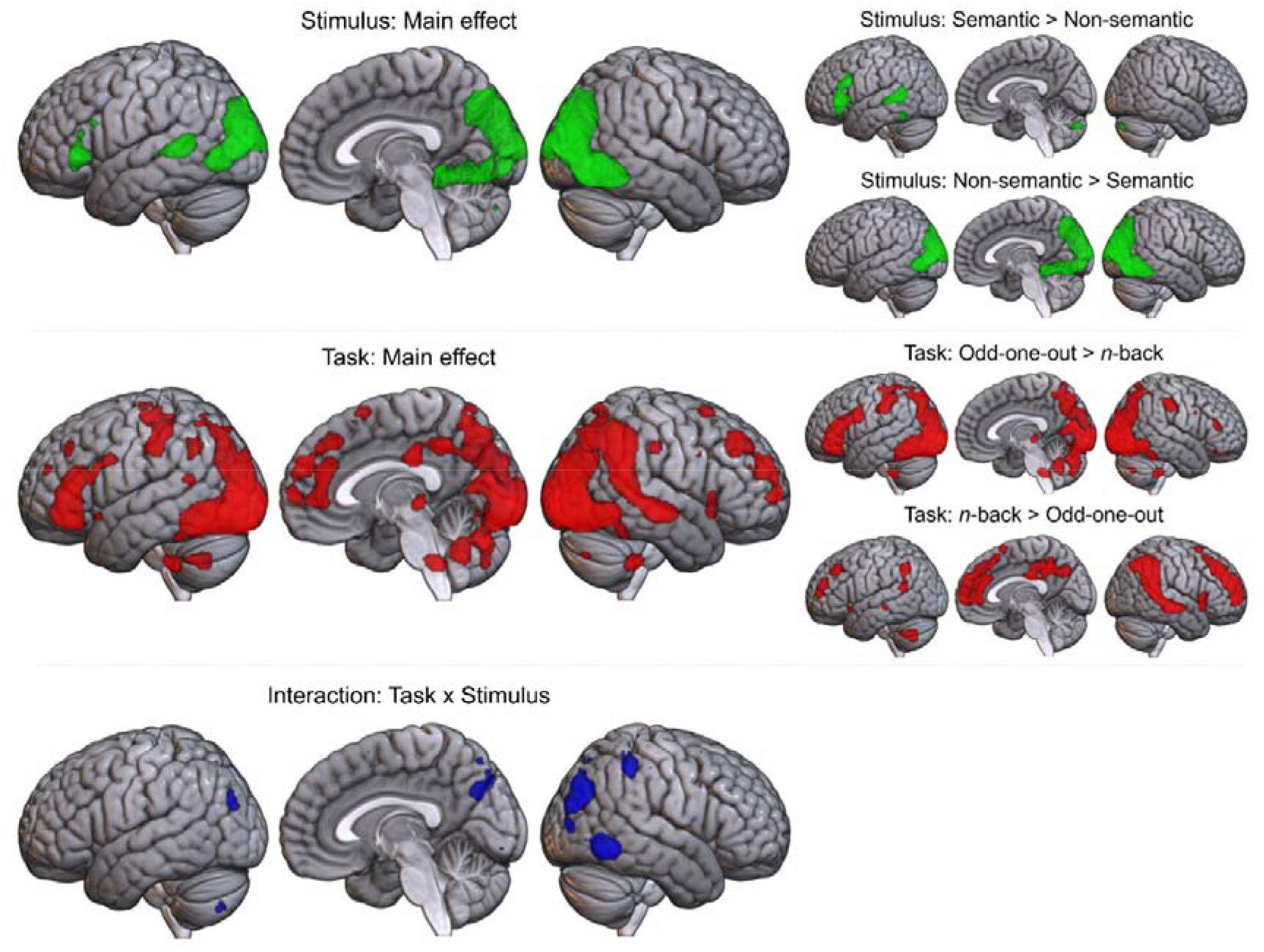
*Results of a two-way ANOVA distinguishing the effects of task process (odd-one-out vs*. n*-back), stimulus domain (semantics vs. non-semantic) and their interaction on the activity associated with hard, controlled processing. Voxel-level threshold p<0*.*05 with FWE correction. Top box: main effect of stimulus (semantic vs. non-semantic). Middle box: main effect of task (odd-one-out vs*. n*-back). Post hoc tests on the right show the direction of both effects. Lower box: regions where activation is affected by the interaction between task process and stimulus domain*.

The main effect of stimulus domain involved bilateral inferior occipital lobe, right posterior ITG/fusiform, left posterior MTG/superior temporal sulcus (STS) and left IFG. Post hoc comparisons showed greater activation in the left IFG (pars triangularis and orbitalis) and left posterior STS/MTG and fusiform gyrus for semantic over non-semantic stimuli. Notably, these clusters are in SCN regions, indicating that there is a significant effect of stimulus domain in areas considered central to semantic control. For non-semantic over semantic stimuli, large clusters were found in left and right inferior and middle occipital gyri. The right hemisphere cluster was larger and extended into posterior fusiform gyrus. The identification of left IFG and pMTG/STS as stimulus-related, even when accounting for the effect of task, confirms the particular importance of these regions for the control of meaningful stimuli.

Post hoc t-tests comparing pairs of the four task-stimulus conditions gave further information about the direction and source of the ANOVA results (see Figure 5 and Supplementary Table 4). Consistent with the SCN template overlap, pairwise comparisons of the semantic odd-one-out condition over either non-semantic odd-one-out or semantic *n*-back conditions, highlight a left-lateralised pattern comprising key SCN regions in left IFG and ventral posterior temporal cortex. This pattern was not found for the semantic > non-semantic *n*-back condition, nor were any other significant effects identified (and the reverse contrast only revealed a small cluster in the inferior occipital lobe). This indicates that the main effects of stimulus were driven largely by differences within the odd-one-out task, while the two variants of the n-back task engaged similar regions, albeit with differences in extent. Both the presence of semantic stimuli and a task requiring the controlled selection and manipulation of this information appear necessary to shift the pattern of activity from domain-general to semantic control areas. This pattern of additive main effects may not hold for the posterior STS/MTG region, which was only found to relate to stimulus domain, although this difference was only identified within the odd-one-out task.

**Figure 5.**
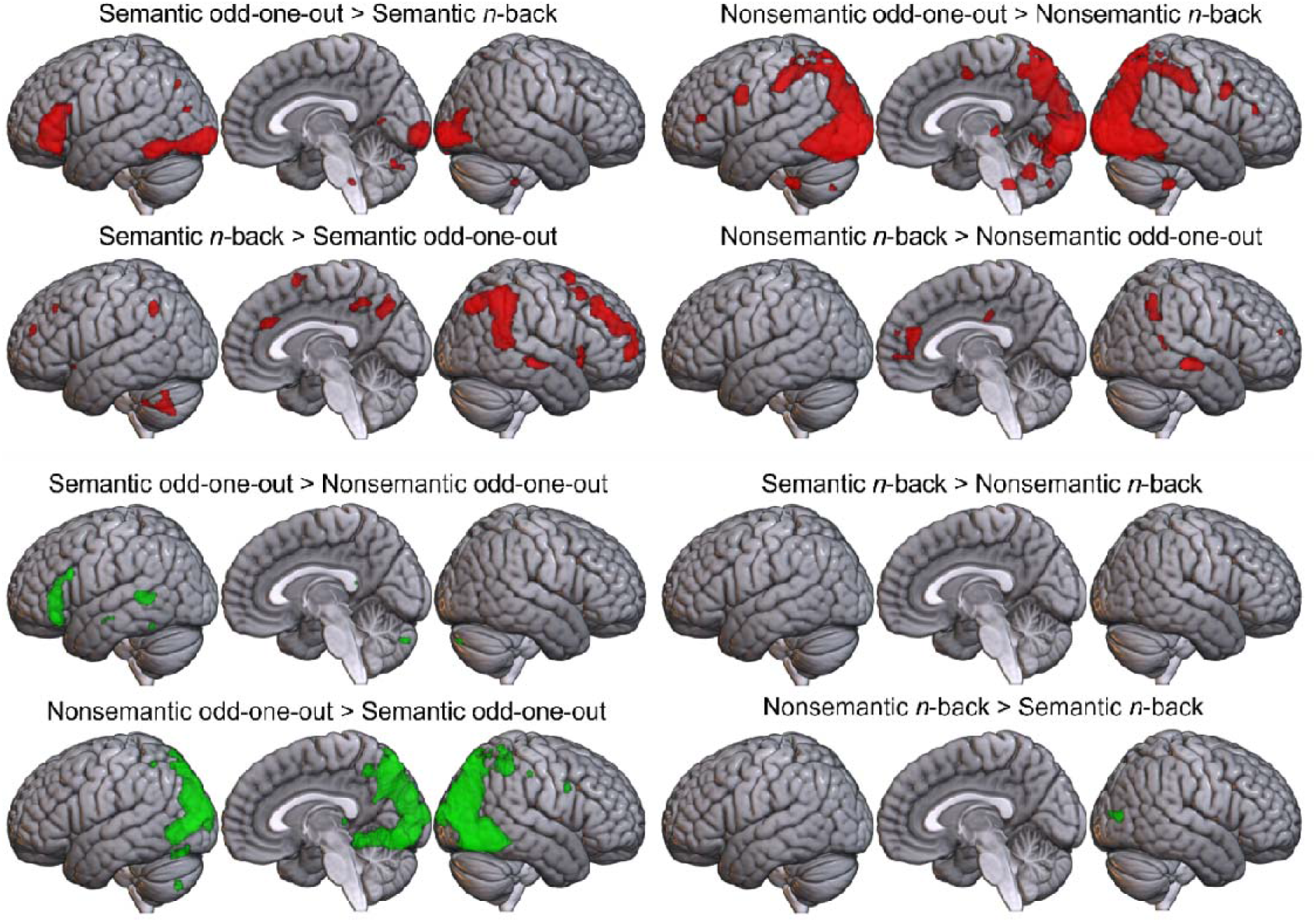
*Pairwise comparisons between conditions. Voxel-level threshold p<0*.*05 with FWE correction. Top left quadrant: semantic odd-one-out vs*. n*-back task. Top right quadrant: non-semantic odd-one-out vs*. n*-back task. Bottom left quadrant: semantic versus non-semantic odd-one-out task. Bottom right quadrant: semantic versus non-semantic* n*-back task. Comparisons assessing the impact of stimulus domain are shown in green, while comparisons assessing task process effects are displayed in red*.

Interaction effects were found in bilateral occipital lobe and right IPL and posterior ITG. These appear to be a result of the large bilateral swathe of occipito-parieto-temporal cortex showing greater activity for non-semantic odd-one-out than both semantic odd-one-out and non-semantic *n*-back conditions. These regions are implicated in the dorsal attention network (DAN), thought to support the top-down control of visual attention ^54–56^, an executive process that may be particularly important for the non-semantic odd-one-out condition given the requirement to selectively attend to different aspects of complex visual stimuli. This may include the planning of eye movements, shown to involve an overlapping region of posterior parietal cortex ^57,58^. These areas drive the higher overlap of the non-semantic odd-one-out than *n*-back condition with the MDN template. However, the areas showing a preference for the *n*-back task regardless of stimulus domain, including bilateral MFG and right IFG, are also core MDN regions. This suggests different subsets of MDN areas are preferentially engaged for different task processes, despite stronger overall overlap with the MDN for the odd-one-out task, while the need to flexibly select and attend to different semantic features engages the SCN.

### ROI analyses

The effects of task process and stimulus domain may vary within and between networks. Individual *a priori* regions of interest (ROIs) within the SCN and MDN were examined using a factorial ANOVA (constructed as per the whole brain analyses) with multiple-comparison corrected post-hoc tests where applicable (all statistics are in Supplementary Table 5). The functional profiles of the ROIs derived from the SCN template (left and right IFG (pars orbitalis and pars triangularis), left pMTG/STS, left pITG and the dmPFC) are shown in Figure 6 (note some of these regions are also identified in the MDN, see below). These SCN ROIs may be split into two broad groups. In the first group, including all left IFG and posterior temporal ROIs, ANOVAs revealed significant interactions or main effects resulting in the greatest activation for the semantic odd-one-out task. In the left IFG (pars triangularis) and left pMTG/STS, significant interaction effects (IFG; F(1,124)=4.19, p<0.05, pMTG/STS; F(1,124)=8.25, p<0.05) were driven by greater effects of stimulus domain in the odd-one-out task. In pITG there was no interaction, but significant main effects of both task process (odd-one-out>n-back, F(1,124)=55.6, p<0.001) and stimulus domain (semantic>non-semantic, F(1,124)=17.1, p<0.001), indicating largely additive effects. In left IFG (pars orbitalis), there was a significant main effect of stimulus only, with greater activation for semantic stimuli in both tasks (F(1,124)=27.5, p<0.001). All these ROIs were significantly engaged in the semantic odd-one-out task compared to rest, while their involvement in other conditions was inconsistent. These subtle differences in activity profile may or may not be meaningful (see Discussion). However, critically, these regions all respond preferentially to the controlled selection and manipulation of semantic stimuli.

**Figure 6.**
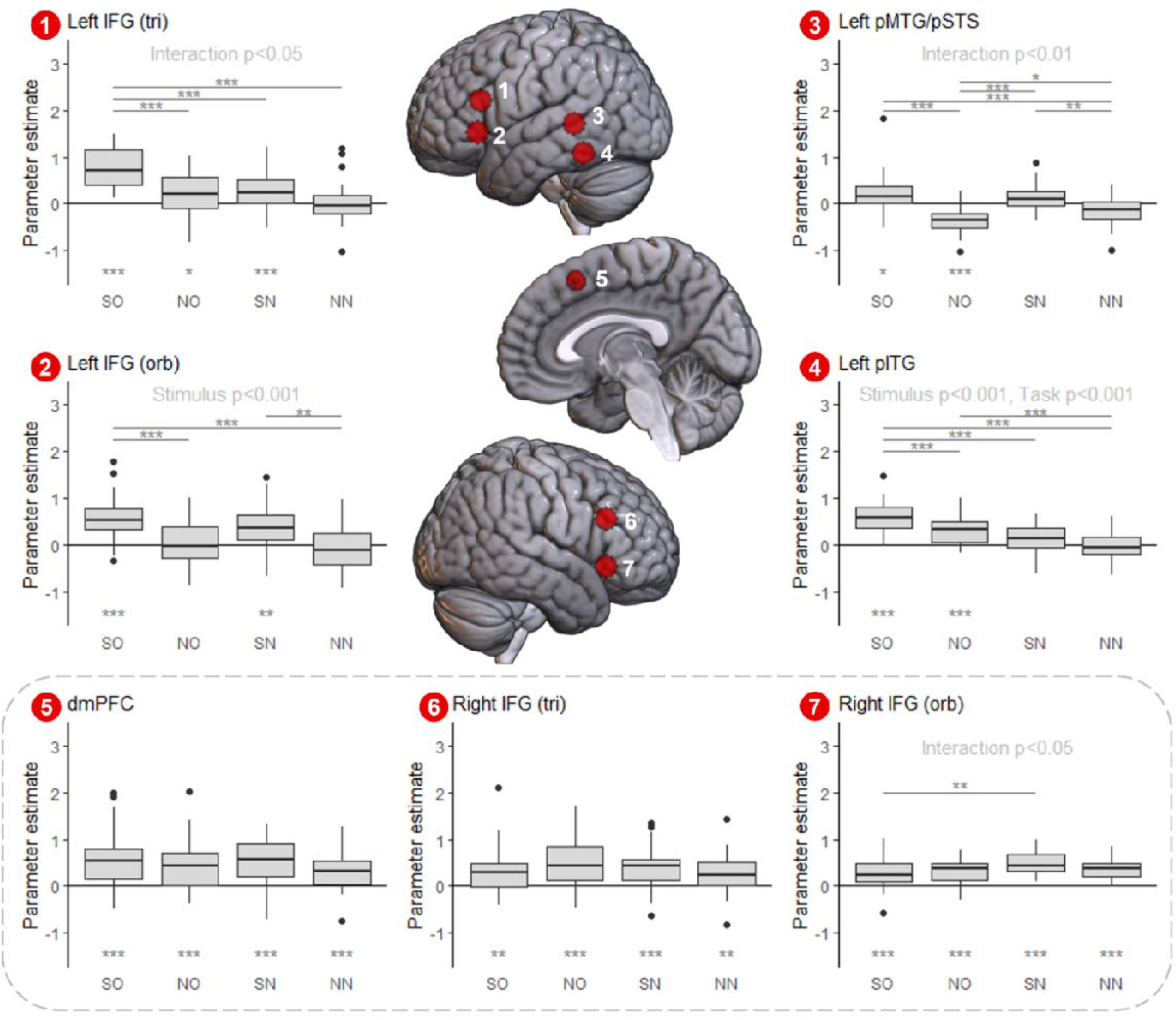
*Functional profile of the ROIs derived from the SCN mask. ROI locations are indicated in the centre panel. Activity for each condition is shown for each ROI (semantic odd-one-out, SO; non-semantic odd-one-out, NO; semantic* n*-back, SN; non-semantic* n*-back, SN). Significant results of two-way ANOVAs (task process x stimulus modality) are shown for each ROI. Multiple comparison corrected post-hoc tests (Tukey’s HSD) between conditions are shown where a significant interaction was observed. One-sample t-tests to determine whether activation is significantly different from rest, are shown with asterisks at the bottom of each graph (multiple comparison corrected within each ROI). Significance levels are indicated with asterisks; *** p<0*.*001, ** p<0*.*01, * p<0*.*05*.

In the second group of ROIs constructed from the SCN mask, the dmPFC and right IFG (pars triangularis and pars orbitalis) were significantly activated across all conditions. The dmPFC and right IFG (pars triangularis) showed no significant interaction or main effects, with consistent activation in each condition. In the right IFG (pars orbitalis), there was a significant interaction effect, F(1,124)=4.50, p<0.05, driven by a difference between the semantic *n-*back and odd-one-out tasks. These regions differ from the other four SCN ROIs, as none show a preference for the semantic stimuli or odd-one-out task. Instead, these regions are activated similarly across conditions, consistent with a role in domain-general control. As the neuroimaging-defined SCN does not exclude regions based on their involvement in control outside of the semantic domain, the identification of some domain-general areas is not unexpected. Here we demonstrate heterogeneity within the *a priori* SCN mask, distinguishing regions preferring semantic control and domain-general areas. This split is highly consistent with prior literature, as only the regions showing a semantic preference overlap the areas where lesions are indicative of semantic aphasia, while the more domain-general areas overlap the MDN.

Figure 7 displays the functional profile of the ROIs derived from the MDN mask (left and right SFG, precentral gyrus (PCG), IPL, anterior MFG (aMFG) and inferior occipito-temporal cortex (IOC)). Note that there would also be MDN ROIs in SMA, insula and mid MFG, but as they show substantial overlap with the SCN ROIs already presented (SMA with dmPFC, bilateral mid MFG with IFG (pars triangularis) and insula for IFG (pars orbitalis)), they are not included in the main text (see Supplementary Figure 4 and Supplementary Tables 5 & 6). A broadly similar pattern of activity was observed across all MDN ROIs except bilateral aMFG: a significant interaction between task and stimulus, which was larger in the right hemisphere. The non-semantic odd-one-out condition showed significantly greater activation than the non-semantic *n*-back condition (in all ROIs), the semantic odd-one-out condition (in bilateral SFG and IOC, right PCG and IPL), and the semantic *n*-back condition (in bilateral PCG and IOC and right IPL), indicating a marked preference for the non-semantic odd-one-out condition across most of the MDN. Despite this preference, most MDN-derived ROIs were consistently activated across all four conditions, unlike the core SCN. However, the bilateral IOC was only strongly engaged by the odd-one-out task.

**Figure 7.**
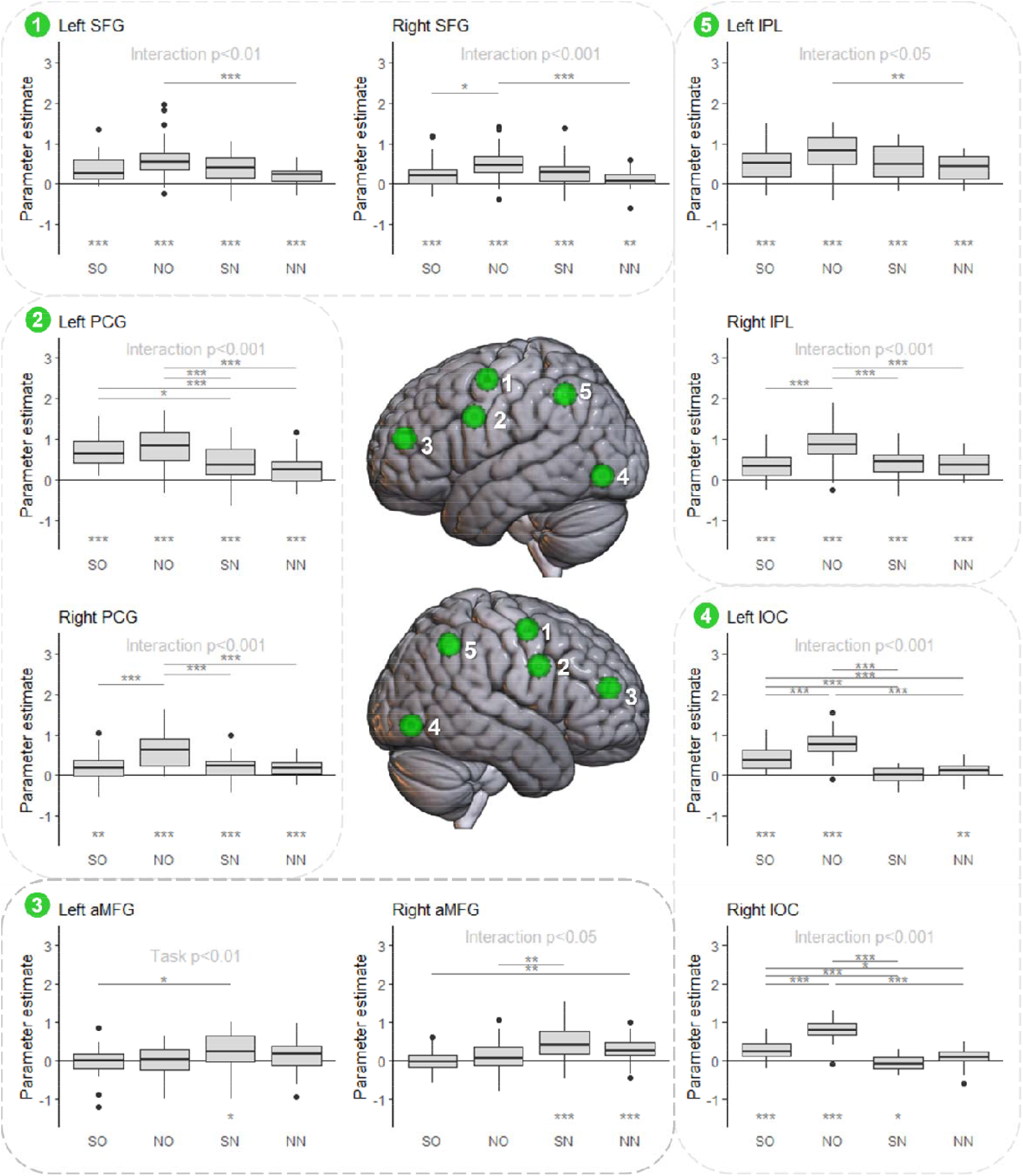
*Functional profile of the ROIs derived from the MDN mask. ROI locations are indicated in the centre panel. Activity for each condition is shown for each ROI (semantic odd-one-out, SO; non-semantic odd-one-out, NO; semantic* n*-back, SN; non-semantic* n*-back, SN). Significant results of two-way ANOVAs (task process x stimulus modality) are shown for each ROI. Multiple comparison corrected post-hoc tests (Tukey’s HSD) between conditions are shown where a significant interaction was observed. One-sample t-tests to determine whether activation is significantly different from rest, are shown with asterisks at the bottom of each graph (multiple comparison corrected within each ROI). Significance levels are indicated with asterisks; *** p<0*.*001, ** p<0*.*01, * p<0*.*05*.

The bilateral aMFG ROIs showed a distinct pattern from the other MDN-derived ROIs: a preference for the *n*-back over the odd-one-out task, reflected in a main effect of task in the left aMFG and an interaction effect in the right aMFG, with significant differences for semantic *n*-back > non-semantic odd-one-out and non-semantic *n*-back > semantic odd-one-out conditions. Neither ROI was significantly activated by either odd-one-out variant relative to rest. Thus, consistent with the whole brain results, the precise pattern of MDN recruitment differs with task process. Activity across the MDN-derived ROIs is therefore not entirely homogenous; while most regions show a preference for the non-semantic odd-one-out condition, the aMFG instead shows an *n*-back task preference regardless of stimulus domain. Furthermore, the dmPFC and right IFG ROIs derived from the SCN mask also overlap the MDN mask and demonstrate equivalent activation across all conditions. Thus, the varying effects of task process and stimulus domain across the MDN may reflect at least three distinct functional profiles.

## DISCUSSION

For the first time, we distinguished the separable effects of task process and the semantic nature of the stimulus on the engagement of control regions across the brain. The key findings were as follows:

1. Both task process and stimulus domain affect the control regions engaged. Previous comparisons of stimulus domain are likely confounded by the strong task effects identified within key regions here. Despite this we demonstrate that differing areas are responsible for semantic and domain-general control even within the same task.
2. The presence of meaningful stimuli is necessary, but not sufficient, to strongly engage the core semantic control regions in left IFG and posterior temporal cortex and shift the activity pattern from resembling the MDN to the SCN. Relevant task processes, such as the searching, selection and inhibition of potential rules, are also required but must be applied to manipulate semantic information.
3. An additional set of regions consistently identified in semantic control tasks (dmPFC and right IFG), was equally active for semantic and non-semantic stimuli and across task processes. This domain- and task-independent involvement is consistent with their additional identification as part of the MDN. However, this pattern was not observed across the rest of the MDN.
4. Instead, the majority of the MDN (bilateral superior frontal, inferior parietal, precentral gyrus and lateral occipital regions) was preferentially engaged for the selection and inhibition of non-semantic stimuli in the odd-one-out task. Similarly, the overall pattern of activation in this condition most closely resembled the MDN.
5. Even excluding regions overlapping the SCN, the MDN was heterogeneous, with bilateral MFG regions displaying greater activation for working memory processes and no stimuli preference.

Disentangling the impact of stimulus domain and task process within the same participants allowed us to test key assumptions in the semantic literature. This study joins a set of emerging evidence that the regions and networks engaged in semantic control are not simply the same as for domain-general control ^35–37,59^. Critically, by directly comparing the semantic and non-semantic odd-one-out variants we demonstrate this effect of stimulus-domain whilst holding task process and difficulty constant. Additionally, we found that the presence of meaningful, semantic stimuli alone is not sufficient to engage the SCN. Instead, the network is preferentially engaged when performing tasks that require the manipulation of meaningful stimuli, via inhibition or selection of semantic features or searching for weak or subordinate meanings ^1,4,5,9^. The working memory and attention processes utilised in an *n*-back task do not require this manipulation, and therefore do not engage the SCN. However, when a task does require this manipulation (as in the odd-one-out task), simply varying the nature of the stimuli results in SCN recruitment. The critical importance of both task and stimulus resonates with the definition of semantic control derived from neuropsychological assessment ^5,12,16,60,61^, while encouraging a more nuanced understanding of its nature and mechanisms.

All regions consistently implicated across studies of semantic control ^4^ were activated for the semantic odd-one-out task. However, their functional profiles were not homogenous, indicating the different roles these areas may play in semantic and non-semantic tasks. A left-lateralised set of SCN regions in IFG and posterior temporal cortex (shown in red in Figure 8C) preferentially activated for semantic stimuli, while the dmPFC and right IFG (shown in yellow) showed a qualitatively different profile of broad domain-generality. Thus, the SCN as previously described may consist of two separable parts: a set of ‘core SCN’ regions with a (relative) specialisation for semantic control in left IFG and posterior temporal cortex, and ‘peripheral SCN’ consisting of domain-general regions in right IFG and dmPFC which may support executive processes that are necessary for, but not unique to, semantic control. The inclusion of some areas without a particular specialisation for semantics within the SCN mask is not surprising, as the network was only defined with reference to semantic cognition. However, the current analyses distinguish the regions with and without this preference. This distinction corresponds extremely well to the neuropsychology data, which has associated semantic control deficits with lesions in the core regions only ^11,12,62^. The peripheral areas may support the same processes when more resources are required or perform a distinct role, for example, determining the corresponding motor action following a semantic decision ^63–65^. These peripheral areas overlap the MDN and could simply be disregarded when considering semantic control. However, controlled semantic cognition did not activate the entire MDN in combination with the SCN. Instead, the functional profile of this small subset of broadly domain-general regions differs from the majority of the MDN which has a particular preference for non-semantic stimuli. This explains why only this subset of MDN regions has been consistently identified in difficult semantic tasks (and conversely why IPL is inconsistently identified) ^4,9^ and suggests a broader support role for these areas working in combination with the rest of the SCN or MDN, as required.

**Figure 8.**
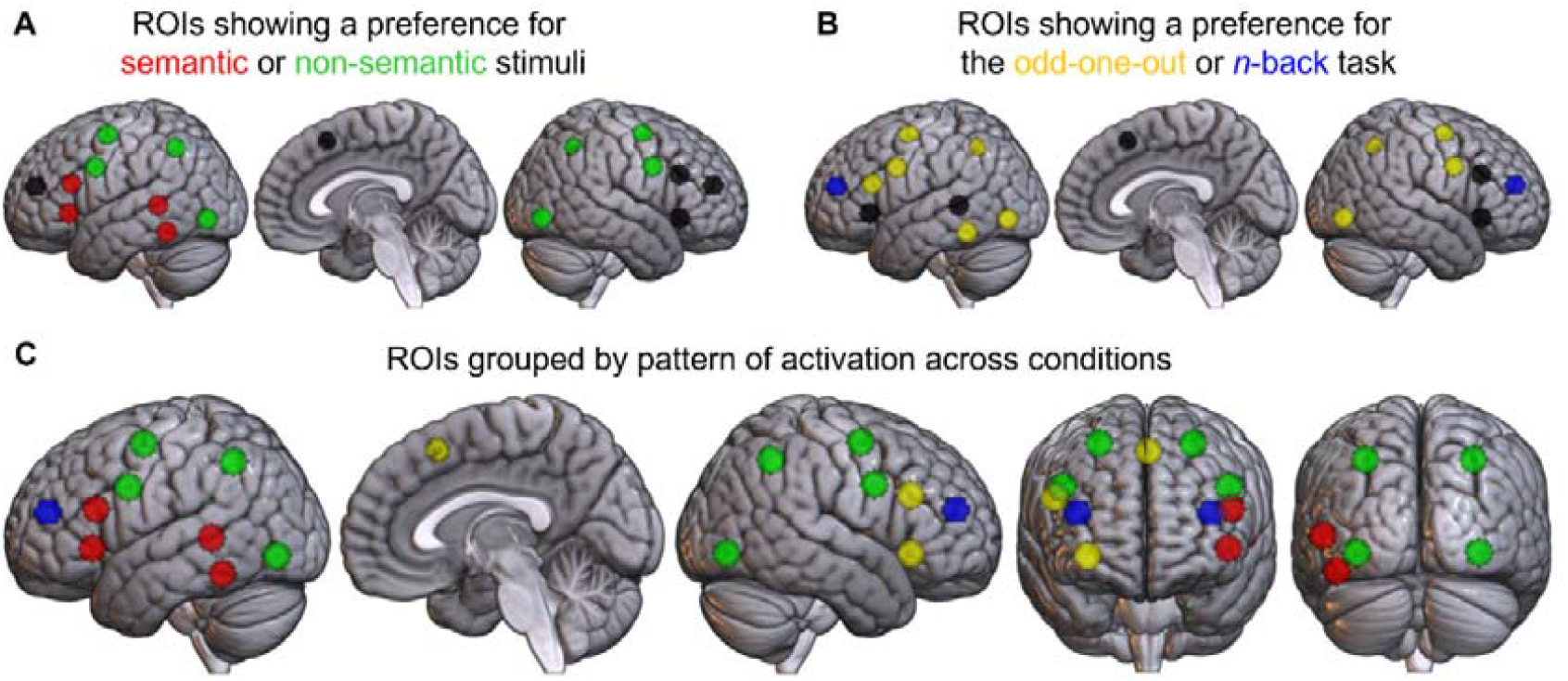
*Summary of activation across ROIs derived from the SCN and MDN masks; results are synthesised from the ANOVA results and post-hoc tests. A) ROIs are coloured by their stimulus domain preferences; those with a preference for semantic stimuli in red, for non-semantic stimuli in green, and for neither in black. B) ROIs are coloured by their task preference; those with a preference for the odd-one-out task in yellow, for the* n*-back task in blue, and for neither in black. C) ROIs are grouped within each network by their functional profile across all conditions. The red SCN regions were preferentially engaged for the manipulation of semantic stimuli, whereas the yellow SCN regions showed similar engagement across tasks and stimuli. The green MDN regions preferentially activated for the non-semantic odd-one-out task, whilst the blue MDN regions were more active for the* n*-back task*.

The functional profile also varied across the remainder of the MDN. The majority of MDN regions (shown in green in Figure 8C) were sensitive to both task and stimuli, preferentially activating for the non-semantic odd-one-out condition. These areas are consistent with prior studies comparing across both task process and stimuli domain simultaneously ^36,42^. In contrast, the bilateral aMFG (shown in blue) showed a preference for the *n*-back task across stimulus domains. This division resembles a separation into the different intrinsic connectivity networks overlapping the functionally-defined MDN. The preference for the non-semantic odd-one-out task appears within the DAN, whilst the other functional groupings overlap the frontoparietal and salience (cingulo-opercular) networks ^66–68^. The inhomogeneous functional profiles across the MDN may help reveal its hidden structure and are consistent with recent proposals that the precise set of MDN regions maximally engaged for a given task shifts based on the paradigm utilised ^33,69^. While large portions of the MDN may support the flexible manipulation of non-semantic information, others like the aMFG (or peripheral SCN) may have a particular role, for instance, in the working memory or attentional engagement processes necessary for the *n*-back task ^45,52,70,71^. The laterality differences in IFG function are also worth noting as MDN results are sometimes summarised across hemispheres ^26,50^.

The preference for non-semantic stimuli found throughout much of the MDN might be surprising given its proposed domain-generality. However, the current study focuses on the semantic domain precisely because this appeared to be an exception to the typical pattern of activation found for executive control across other domains. The direct comparison of the semantic and non-semantic odd-one-out variants demonstrates the presence of this stimulus domain difference even when task process, difficulty and time-on-task are held constant. Thus, there may be something ‘special’ about the manipulation of semantic information which requires the recruitment of the SCN as a specialised resource, that cannot be resolved by recruiting the more domain-general MDN. It is not yet known whether the semantic domain is the only exception, or why this might be the case. Semantic control may rely on different regions due to the need to select and manipulate internal, stored information, as opposed to information present within an environmental stimulus. This would point to episodic memory as another domain which could rely on the same regions as semantic control, a hypothesis for which neuropsychological and neuroimaging studies have shown initial support ^72,73^. Alternatively, these results may not be driven solely by the meaningful nature of the stimuli. In the present work, we have maximised our ability to detect the stimulus domain effects by contrasting meaningful verbal with meaningless visuospatial stimuli, as typically used to study each domain. Thus, these stimulus effects may, at least in part, relate to the visuospatial or verbal nature of the stimuli. Further research should distinguish the effects of varying different aspects of the stimulus. Additionally, as the different control regions display diverse profiles even when only studying two domains, it may be necessary to move beyond a simple dichotomy of domain-specific versus domain-general control ^74,75^.

These findings have implications for the functional organisation of frontal and posterior temporal control areas. Nearby frontal regions display distinct functional profiles. Both left pars triangularis and pars orbitalis show a preference for semantic stimuli, while left PCG demonstrates the opposite pattern. All these areas prefer the odd-one-out condition, while aMFG shows greater activation for the *n*-back task, and no domain preference. Thus, the core left IFG semantic control region appears bordered by domain-general control regions with distinct roles, consistent with prior comparisons between coarse language and control networks ^34,76^. This pattern of nearby, interdigitated networks does not appear to be mirrored elsewhere, with parietal cortex demonstrating a non-semantic preference only. In left posterior temporal cortex, prior work implicated a broad region including both pMTG/STS and pITG ^4^ in semantic control, and a pITG region in the extended MDN ^33^. This led to the suggestion of distinct superior and inferior regions for domain-specific and domain-general control, respectively ^4,75,77^. Yet, here, both pITG and pMTG/STS show a preference for the manipulation of semantic stimuli, forming part of the core SCN. However, pMTG/STS is significantly deactivated to the non-semantic odd-one-out condition, while pITG is significantly activated, suggesting that both have a relative preference for semantic stimuli, but pITG does not exclusively support semantic control. Alternatively, distinct pITG neuronal ensembles may support control over the two domains and only appear to overlap.

The main analyses presented here used rest and not the easy task variant as a baseline for two reasons: first, to avoid a confound of difficulty when comparing the semantic and non-semantic odd-one-out task variants which would compromise the ability to distinguish the effects of stimuli domain and task process. Second, both the hard and easy odd-one-out conditions over rest strongly engage control areas, suggesting the easy variant is too difficult to serve as an appropriate baseline as key control regions would be missed. The hard > easy analyses in the Supplementary Materials demonstrate both issues. While it is preferable to use an active baseline to factor out regions responsible for the representation of the stimuli, the areas identified here are known to support control processes based on their strong similarity with the *a priori* templates and their identification in the hard>easy contrasts. This includes the posterior DAN areas which may relate to the control of visuospatial stimuli or a greater need for controlled eye movements. While we were able to rule out difficulty effects in the stimulus domain, the diverse nature of our tasks made matching their difficulty impossible. However, they engaged qualitatively different networks of regions.

These results demonstrate the influence of multiple factors on the recruitment of control regions and the need to carefully disentangle each factor. Core semantic control regions preferentially activate for meaningful stimuli, even when task processes remain constant. In contrast, most MDN regions perform similar task processes, yet demonstrate a non-semantic preference. Both sets of areas may be supported by the peripheral SCN regions implicated in control regardless of domain. Why meaningful stimuli require different control areas is a critical avenue for further research.

## METHODS

### Participants

All participants were right-handed native English speakers, with no brain injury or learning disorders. They provided written informed consent and received monetary compensation. Of the 40 participants scanned, 3 were excluded for excessive head motion (>3mm translation or >1° rotation in any run). A further 5 were excluded for poor behavioural performance. These participants did not follow instructions to prioritise accuracy over speed in the odd-one-out tasks and subsequently finished the trials in one or more runs early, resulting in insufficient imaging data. 3 such participants also showed lower accuracy scores for the easy than the hard condition in at least one task variant, indicating a guessing tactic. The final sample consisted of 32 neurotypical participants (achieving the desired sample size determined by *a priori* power analysis in G*Power 3.1 ^78^, desired □ = 0.8 and a medium effect size, Cohen’s d = 0.5) aged between 18-28 (mean 21.0 ± 3.08 years; 17 females, 15 males).

### Tasks and materials

Two tasks were used, the odd-one-out task and the *n-*back task. Two variants of each task were constructed, one using meaningful semantic stimuli (words), and the other using meaningless nonverbal stimuli (greyscale geometric shapes). This 2x2 design allowed comparison of semantic to non-semantic stimuli, while independently varying the cognitive processes used. Semantic stimuli were verbal, and non-semantic stimuli nonverbal, to maximise differences between stimulus types and remain consistent with the majority of prior assessments of semantic control ^4^ and domain-general control ^26,28,31,33,40^. Each variant was performed at two levels of difficulty (hard and easy). An independent sample of 14 native English speakers took part in a behavioural pilot to attempt to match RT and accuracy of the conditions at each difficulty level. All task items are included in the Supplementary Materials.

#### Semantic stimuli

Semantic stimuli comprised meaningful, concrete English words, 3-10 letters long, taken from the N-watch word list ^79^. Imageability, frequency and age of acquisition were matched across conditions (p > 0.05). All words used in the hard variant of the odd-one-out task were shuffled and recombined to create the easy variant, improving the matching of stimuli characteristics across conditions, with additional words included as more easy trials were needed. The semantic *n*-back task used the same words as the easy semantic odd-one-out task.

#### Non-semantic stimuli

The non-semantic stimuli were greyscale, meaningless geometric patterns designed to have no verbal or semantic content, bounded within a square, in the style of the Cattell Culture Fair task. Following Woolgar et al. ^28^ there were 4 items per trial instead of 5 for greater suitability for neuroimaging. 84 individual items, comprising 21 hard trials, were taken from Woolgar et al. ^28^. The three additional trials that contained semantic content were not used. The remainder of the items were created for the present study following the same style.

Where possible (i.e., where there was sufficient dissimilarity), items in the hard odd-one-out trials were also used in the easy variant of the odd-one-out task, to increase the visual similarity between conditions. All items from the easy odd-one-out variant were used to create all trials of the *n*-back task, as with the semantic variant.

#### Odd-one-out task design

The non-semantic variant of the odd-one-out task was initially adapted from the Cattell Culture Fair test battery by Woolgar et al. ^28^ and further adapted here (described in detail above). A semantic variant of this task was created using single word stimuli. In both variants, participants determined which item was the ‘odd-one-out’ and pressed the corresponding button on each trial. Items were displayed for the entire trial duration. Participants were encouraged not to guess, but to respond when they were sure of the rule. The trial ended either after a response was made, or after 20 seconds to ensure that participants did not disengage from the task if they were unable to solve a given trial. The odd-one-out task was therefore self-paced following Woolgar et al. ^28^, to maximise time on-task and match this across conditions (as time-on-task would differ between hard and easy conditions if trials had a fixed duration).

At the easy difficulty level, the target item is simple to detect as the most salient feature is indicative of the correct rule, e.g., alligator is the odd-one-out in table/sofa/alligator/cabinet, as it is the only living item, and the large square is the odd-one-out in small triangle/large square/small circle/small hexagon, as it is the only large shape. In the harder conditions the rule needed to identify a single target item is harder to detect to promote searching through different possible rules, inhibition of dominant features associated with incorrect rules, and rule-switching. In some hard trials, the rule is simply harder to generate or detect (e.g., ‘mallet’ is the odd-one-out in jewel/yacht/mansion/mallet, as the only non-expensive item; two triangles is the odd-one-out in five pentagons/two triangles/four squares/six hexagons as the number of shapes must match the number of sides). In others, some items initially suggest an incorrect rule that needs to be suppressed (e.g., ‘branch’ is the odd-one-out in branch/wing/cabin/engine as it isn’t part of a plane, but ‘branch’ and ‘wing’ both share an association with birds; the large anticlockwise spiral is the odd-one-out in large clockwise spiral/small clockwise spiral/large anticlockwise spiral/small clockwise spiral as size is irrelevant but salient and would typically be noticed before spiral direction). Thus, the semantic and non-semantic versions of the odd-one-out task are designed to strongly engage the same selection, inhibition and rule searching processes, considered central to both semantic and domain-general control, differing only in the stimulus domain in which these processes are performed. Each task variant is shown in Figure 1 and all trials are presented in the Supplementary Materials.

#### n*-back task design*

A standard *n*-back task design was used (see Figure 1). In each block, 15 items were presented sequentially. Participants were required to attend to the sequence and press a button if the current item matched the item presented *n* items ago. Following previous *n*-back paradigms ^70,71^, each item was displayed for 500ms, with an interstimulus interval of 1500ms and a total trial length of 2 seconds. Participants completed a 3-back (hard) and a 1-back (easy) variant of the task for each stimulus type. In the semantic variant, words were presented in upper or lower case in a randomised manner, to decrease the use of the visual word form to solve the task. Note however that task performance does not necessitate deep semantic processing as participants may use other strategies, such as phonological working memory. Here this condition is labelled ‘semantic *n*-back’ to highlight the presence of semantic stimuli, not to specify the process used. Indeed, the key question asked here is whether the presence of meaningful semantic stimuli alone is sufficient to change the pattern of control-related activation, without further changes in task process.

### Experimental design

Participants took part in a single scanning session, comprised of a structural scan and four functional runs. Participants completed each of the four task-stimulus conditions in a separate run. Within each run, participants alternated between hard and easy blocks with two rest blocks at set positions (one third and two thirds through). This was necessary to allow each hard condition of interest to be compared to its respective easy condition and the rest baseline with sufficient power. If all 9 conditions were evenly spaced across the four runs, a standard high pass filter would disallow the critical comparisons of interest. Each block lasted 31 seconds, for a total run length of 806 seconds. The order of runs and whether the first block of each run was hard or easy was counterbalanced across participants.

### fMRI data acquisition

Imaging data were acquired using a 3T Siemens PRISMA scanner with a 32-channel headcoil. T1-weighted anatomical images were acquired using a 3D Magnetisation Prepared Rapid Gradient Echo (MPRAGE) sequence, with the following parameters: repetition time (TR) = 2.25s, echo time (TE) = 3.02ms, inversion time = 900ms, flip angle = 9°, field of view (FOV) = 256 x 256 x 192mm, GRAPPA acceleration factor 2, 1mm resolution.

Functional images were acquired using a multi-echo multi-band sequence, to promote signal across the entire brain, including key semantic regions where signal loss and distortion is typical, such as the anterior temporal lobe ^80,81^. The sequence had the following parameters: TR = 1.792s, four echoes with echo times TE_1_ = 13ms, TE_2_ = 25.85ms, TE_3_ = 38.7ms, TE_4_ = 51.55ms, flip angle 75°, FOV = 240 x 240 x 138mm, multi-band factor acceleration factor 2, GRAPPA acceleration factor 2. Each EPI volume consisted of 46 axial slices with resolution of 3mm. For each participant, 1800 volumes were acquired in total across four runs, each lasting 806 seconds.

### fMRI pre-processing

Raw DICOM files were converted to BIDS-compatible NIfTI files using HeuDiConv from Nipype ^82^. The fMRIprep pipeline was then used to preprocess all anatomical and functional data ^83^. For anatomical data, the pipeline consisted of four steps. First, the T1-weighted image for each participant was corrected for intensity non-uniformity correction using N4BiasFieldCorrection ^84^ distributed with ANTS ^85^. The corrected T1-weighted image was skull-stripped using the antsBrainExtraction workflow with OASIS30ANTs as a target template. Brain tissue segmentation of cerebrospinal fluid, white matter and grey matter was then performed using FAST (FSL 5.0.9, ^86^). Finally, images were normalised to MNI space using nonlinear registration with antsRegistration (ANTS 2.3.3 ^85^) and the MNI152Lin2009cAsym template.

For each run of functional data, a reference volume and its skull-stripped version were generated from the shortest echo. This BOLD reference was then co-registered to the T1-weighted reference using FLIRT (FSL 5.0.9, ^87^), and head-motion parameters with respect to the BOLD reference were estimated using MCFLIRT (FSL 5.0.9, ^88^). BOLD runs were then slice-time corrected using 3dTshift from AFNI ^89^, and the corrected BOLD time-series were resampled into native space by applying the transforms to correct for head motion. The four echo-times were then optimally combined using a weighted T2* map following the method by Posse et al. ^90^, calculated using a monoexponential signal decay model with nonlinear regression. These optimally-combined BOLD time-series were resampled into standard MNI space and smoothed using an 8mm FWHM Gaussian kernel in SPM12 in MATLAB R2018a.

### Whole brain analysis

Using SPM12 (https://www.fil.ion.ucl.ac.uk/spm/software/spm12/) in MATLAB R2018a, data were high-pass filtered at a cut-off of 128s and then analysed using a general linear model (GLM). The factorial experimental design featured two tasks and two stimulus modalities, yielding four task-stimulus conditions, each of which were performed at two levels of difficulty for a total of eight experimental conditions. Rest was included as an implicit ninth condition in the GLM, across runs as a single regressor. At the first level, the estimated head motion parameters and run means were included as covariates of no interest. The hard and easy variants of each condition were each contrasted over rest, and the hard variant contrasted over the easy variant. Each of these contrasts was then carried forward in to the second level and statistically thresholded at an FWE-corrected voxel level of p<0.05.

To assess whether the activity pattern for each task variant resembled existing assessments of the MDN or SCN and whether varying task process or stimulus modality drives a transition between these typical semantic and domain-general control patterns, the hard>rest results for each variant were compared to templates of the SCN and MDN. Template masks were acquired for each network; the MDN template was taken from Fedorenko et al. ^26^ (thresholded at t>1.5 following the authors) and the SCN template was the result of Jackson’s meta-analysis ^4^ (thresholded at the voxel level p<0.001 and at the cluster level with family-wise error correction p<0.001 following the author). The Jaccard similarity index ^91^ between the SCN and MDN templates and the thresholded group-level activity was calculated for each of the four task-stimulus variants (in the hard condition > rest). This means the intersection of the voxels active in both the template and the activation map was divided by the union of voxels active in either image.

To assess the impact of the task and stimulus factors, a factorial ANOVA was constructed at the second level, followed by pairwise post-hoc t-tests to determine the direction of effects. The ANOVA utilised the first-level hard>rest contrast images for each of the four conditions. This was considered a fairer comparison than the hard>easy images, after reviewing the full pattern of behavioural and imaging data for each task and stimulus pair (see Results for a more detailed discussion). However, see Supplementary Figures 2 & 3 for similar results utilising the hard>easy images. All group analyses were thresholded at an FWE-corrected voxel level of p<0.05.

### ROI analysis

To investigate the functional profiles of different control regions, analyses were conducted within *a priori* ROIs, using the two thresholded templates as established definitions of each network (i.e., the Jackson SCN meta-analysis ^4^ and the Fedorenko et al. ^26^ MDN mask). 8mm spherical ROIs were constructed around the strongest peak for each cluster using MarsBaR ^92^. Additional peaks were taken from large clusters if those peaks crossed into different anatomical regions. There were seven SCN ROIs: left and right IFG (pars triangularis), left and right IFG (pars orbitalis), left pMTG/pSTS, left pITG and dmPFC. There were also fifteen MDN ROIs: left and right SFG, PCG, MFG, anterior MFG, IPL, inferior occipital cortex and insula, and SMA. However, due to overlap with the SCN ROIs, some of these MDN-derived ROIs are only shown in Supplementary Figure 4. This includes the bilateral MFG ROIs (overlapping with bilateral IFG (pars triangularis)), SMA ROI (overlapping the dmPFC ROI) and the right insula ROI (overlapping the IFG (pars orbitalis)). All peaks used to define the ROIs are in Supplementary Table 6. Contrast estimates were extracted at each ROI (averaged across voxels), for each participant, in each task-stimulus condition (for the hard blocks > rest). One-sample t-tests, multiple comparison corrected within each ROI, were used to determine whether activation in each condition differed significantly from rest. Factorial ANOVAs (task x stimulus) were followed by multiple comparison-corrected pairwise post hoc tests (Tukey’s HSD) where a significant interaction was found.

## Acknowledgements

The authors would like to thank Dr Ajay Halai for his advice and the use of his code for the purposes of data pre-processing, and Dr Alexandra Woolgar for sharing her stimuli for the non-semantic odd-one-out task.

## Notes

**Funding:** This work was supported by a Biotechnology and Biological Sciences Research Council studentship to V.J.H, a programme grant to M.A.L.R. from the Medical Research Council (grant no. MR/R023883/1), an Advanced Grant from the European Research Council to M.A.L.R. (GAP: 670428) and Medical Research Council intramural funding (no. MC_UU_00005/18).

**Data and Code Availability:** The SPM12 toolboxes and fMRIprep pipeline are freely available for download (https://www.fil.ion.ucl.ac.uk/spm/software/spm12/, https://fmriprep.org/en/stable/). All scripts used, task items and results files can be found on GitHub (https://github.com/Vicki-H/Disentangling-task-stimulus), and all task items can also be found in the Supplementary Materials. Raw imaging data can be downloaded from the MRC Cognition and Brain Sciences Unit (https://www.mrc-cbu.cam.ac.uk/publications/opendata/).

### Competing Interest Statement

The authors have declared no competing interest.

## REFERENCES

1. Lambon Ralph, M. A., Jefferies, E., Patterson, K. & Rogers, T. T. The neural and computational bases of semantic cognition. Nat. Rev. Neurosci. 18, 42–55 (2017).

2. Chiou, R., Humphreys, G. F., Jung, J. & Lambon Ralph, M. A. Controlled semantic cognition relies upon dynamic and flexible interactions between the executive ‘semantic control’ and hub-and-spoke ‘semantic representation’ systems. Cortex 103, 100–116 (2018).

3. Patterson, K. & Lambon Ralph, M. A. Chapter 61 - The Hub-and-Spoke Hypothesis of Semantic Memory. in Neurobiology of Language (eds. Hickok, G. & Small, S. L.) 765–775 (Academic Press, 2016). doi:10.1016/B978-0-12-407794-2.00061-4.

4. Jackson, R. L. The neural correlates of semantic control revisited. NeuroImage 224, 117444 (2020).

5. Jefferies, E. The neural basis of semantic cognition: Converging evidence from neuropsychology, neuroimaging and TMS. Cortex 49, 611–625 (2013).

6. Badre, D., Poldrack, R. A., Paré-Blagoev, E. J., Insler, R. Z. & Wagner, A. D. Dissociable Controlled Retrieval and Generalized Selection Mechanisms in Ventrolateral Prefrontal Cortex. Neuron 47, 907–918 (2005).

7. Hoffman, P., Jefferies, E. & Lambon Ralph, M. A. Ventrolateral Prefrontal Cortex Plays an Executive Regulation Role in Comprehension of Abstract Words: Convergent Neuropsychological and Repetitive TMS Evidence. J. Neurosci. 30, 15450–15456 (2010).

8. Davey, J. et al. Exploring the role of the posterior middle temporal gyrus in semantic cognition: Integration of anterior temporal lobe with executive processes. NeuroImage 137, 165–177 (2016).

9. Noonan, K. A., Jefferies, E., Visser, M. & Lambon Ralph, M. A. Going beyond Inferior Prefrontal Involvement in Semantic Control: Evidence for the Additional Contribution of Dorsal Angular Gyrus and Posterior Middle Temporal Cortex. J. Cogn. Neurosci. 25, 1824–1850 (2013).

10. Hoffman, P., Binney, R. J. & Lambon Ralph, M. A. Differing contributions of inferior prefrontal and anterior temporal cortex to concrete and abstract conceptual knowledge. Cortex 63, 250–266 (2015).

11. Noonan, K. A., Jefferies, E., Corbett, F. & Lambon Ralph, M. A. Elucidating the Nature of Deregulated Semantic Cognition in Semantic Aphasia: Evidence for the Roles of Prefrontal and Temporo-parietal Cortices. J. Cogn. Neurosci. 22, 1597–1613 (2009).

12. Jefferies, E. & Lambon Ralph, M. A. Semantic impairment in stroke aphasia versus semantic dementia: a case-series comparison. Brain 129, 2132–2147 (2006).

13. Jefferies, E., Patterson, K. & Lambon Ralph, M. A. Deficits of knowledge versus executive control in semantic cognition: Insights from cued naming. Neuropsychologia 46, 649–658 (2008).

14. Mummery, C. J. et al. A voxel-based morphometry study of semantic dementia: Relationship between temporal lobe atrophy and semantic memory. Ann. Neurol. (2000) doi:10.1002/1531-8249(200001)47:1<36::AID-ANA8>3.0.CO;2-L.

15. Bozeat, S., Lambon Ralph, M. A., Patterson, K., Garrard, P. & Hodges, J. R. Non-verbal semantic impairment in semantic dementia. Neuropsychologia 38, 1207–1215 (2000).

16. Corbett, F., Jefferies, E., Ehsan, S. & Lambon Ralph, M. A. Different impairments of semantic cognition in semantic dementia and semantic aphasia: evidence from the non-verbal domain. Brain 132, 2593–2608 (2009).

17. Kramer, J. H. et al. Distinctive Neuropsychological Patterns in Frontotemporal Dementia, Semantic Dementia, And Alzheimer Disease. Cogn. Behav. Neurol. 16, 211–218 (2003).

18. Pengas, G. et al. Lost and Found: Bespoke Memory Testing for Alzheimer’s Disease and Semantic Dementia. J. Alzheimers Dis. 21, 1347–1365 (2010).

19. Jefferies, E., Patterson, K., Jones, R. W., Bateman, D. & Lambon Ralph, M. A. A category-specific advantage for numbers in verbal short-term memory: Evidence from semantic dementia. Neuropsychologia 42, 639–660 (2004).

20. Whitney, C., Kirk, M., O’Sullivan, J., Lambon Ralph, M. A. & Jefferies, E. The Neural Organization of Semantic Control: TMS Evidence for a Distributed Network in Left Inferior Frontal and Posterior Middle Temporal Gyrus. Cereb. Cortex 21, 1066–1075 (2011).

21. Davey, J. et al. Automatic and Controlled Semantic Retrieval: TMS Reveals Distinct Contributions of Posterior Middle Temporal Gyrus and Angular Gyrus. J. Neurosci. 35, 15230–15239 (2015).

22. Krieger-Redwood, K. & Jefferies, E. TMS interferes with lexical-semantic retrieval in left inferior frontal gyrus and posterior middle temporal gyrus: Evidence from cyclical picture naming. Neuropsychologia 64, 24–32 (2014).

23. Hoffman, P., Pobric, G., Drakesmith, M. & Lambon Ralph, M. A. Posterior middle temporal gyrus is involved in verbal and non-verbal semantic cognition: Evidence from rTMS. Aphasiology 26, 1119–1130 (2012).

24. Duncan, J. The multiple-demand (MD) system of the primate brain: mental programs for intelligent behaviour. Trends Cogn. Sci. 14, 172–179 (2010).

25. Duncan, J. The Structure of Cognition: Attentional Episodes in Mind and Brain. Neuron 80, 35–50 (2013).

26. Fedorenko, E., Duncan, J. & Kanwisher, N. Broad domain generality in focal regions of frontal and parietal cortex. Proc. Natl. Acad. Sci. 110, 16616–16621 (2013).

27. Woolgar, A., Duncan, J., Manes, F. & Fedorenko, E. Fluid intelligence is supported by the multiple-demand system not the language system. Nat. Hum. Behav. 2, 200–204 (2018).

28. Woolgar, A., Bor, D. & Duncan, J. Global Increase in Task-related Fronto-parietal Activity after Focal Frontal Lobe Lesion. J. Cogn. Neurosci. 25, 1542–1552 (2013).

29. Jackson, J., Rich, A. N., Williams, M. A. & Woolgar, A. Feature-selective Attention in Frontoparietal Cortex: Multivoxel Codes Adjust to Prioritize Task-relevant Information. J. Cogn. Neurosci. 29, 310–321 (2017).

30. Assem, M., Glasser, M. F., Essen, D. C. V. & Duncan, J. A Domain-general Cognitive Core defined in Multimodally Parcellated Human Cortex. bioRxiv 517599 (2019) doi:10.1101/517599.

31. Camilleri, J. A. et al. Definition and characterization of an extended multiple-demand network. NeuroImage 165, 138–147 (2018).

32. Müller, V. I., Langner, R., Cieslik, E. C., Rottschy, C. & Eickhoff, S. B. Interindividual differences in cognitive flexibility: influence of gray matter volume, functional connectivity and trait impulsivity. Brain Struct. Funct. 220, 2401–2414 (2015).

33. Assem, M., Glasser, M. F., Van Essen, D. C. & Duncan, J. A Domain-General Cognitive Core Defined in Multimodally Parcellated Human Cortex. Cereb. Cortex 30, 4361–4380 (2020).

34. Fedorenko, E., Duncan, J. & Kanwisher, N. Language-Selective and Domain-General Regions Lie Side by Side within Broca’s Area. Curr. Biol. 22, 2059–2062 (2012).

35. Chiou, R., Jefferies, E., Duncan, J., Humphreys, G. F. & Lambon Ralph, M. A. A middle ground where executive control meets semantics: the neural substrates of semantic control are topographically sandwiched between the multiple-demand and default-mode systems. Cereb. Cortex N. Y. N bhac358 (2022) doi:10.1093/cercor/bhac358.

36. Humphreys, G. F. & Lambon Ralph, M. A. Mapping Domain-Selective and Counterpointed Domain-General Higher Cognitive Functions in the Lateral Parietal Cortex: Evidence from fMRI Comparisons of Difficulty-Varying Semantic Versus Visuo-Spatial Tasks, and Functional Connectivity Analyses. Cereb. Cortex 27, 4199–4212 (2017).

37. Gao, Z. et al. Distinct and common neural coding of semantic and non-semantic control demands. NeuroImage 236, 118230 (2021).

38. Jackson, J. B., Feredoes, E., Rich, A. N., Lindner, M. & Woolgar, A. Concurrent neuroimaging and neurostimulation reveals a causal role for dlPFC in coding of task-relevant information. Commun. Biol. 4, 1–16 (2021).

39. Crittenden, B. M. & Duncan, J. Task Difficulty Manipulation Reveals Multiple Demand Activity but no Frontal Lobe Hierarchy. Cereb. Cortex 24, 532–540 (2014).

40. Duncan, J. & Owen, A. M. Common regions of the human frontal lobe recruited by diverse cognitive demands. Trends Neurosci. 23, 475–483 (2000).

41. Jackson, R. L., Hoffman, P., Pobric, G. & Lambon Ralph, M. A. The Nature and Neural Correlates of Semantic Association versus Conceptual Similarity. Cereb. Cortex 25, 4319–4333 (2015).

42. Krieger-Redwood, K., Teige, C., Davey, J., Hymers, M. & Jefferies, E. Conceptual control across modalities: graded specialisation for pictures and words in inferior frontal and posterior temporal cortex. Neuropsychologia 76, 92–107 (2015).

43. Wang, X., Gao, Z., Smallwood, J. & Jefferies, E. Both Default and Multiple-Demand Regions Represent Semantic Goal Information. J. Neurosci. 41, 3679–3691 (2021).

44. Cattell, R. B. Abilities: their structure, growth, and action. xxii, 583 (Houghton Mifflin, 1971).

45. Owen, A. M., McMillan, K. M., Laird, A. R. & Bullmore, E. N-back working memory paradigm: A meta-analysis of normative functional neuroimaging studies. Hum. Brain Mapp. 25, 46–59 (2005).

46. Gevins, A. & Cutillo, B. Spatiotemporal dynamics of component processes in human working memory. Electroencephalogr. Clin. Neurophysiol. 87, 128–143 (1993).

47. Diamond, A. Executive Functions. Annu. Rev. Psychol. 64, 135–168 (2013).

48. Baddeley, A. D. Working memory. Curr. Biol. 20, R136–R140 (2010).

49. Cole, M. W. & Schneider, W. The cognitive control network: Integrated cortical regions with dissociable functions. NeuroImage 37, 343–360 (2007).

50. Assem, M., Blank, I. A., Mineroff, Z., Ademoğlu, A. & Fedorenko, E. Activity in the fronto-parietal multiple-demand network is robustly associated with individual differences in working memory and fluid intelligence. Cortex 131, 1–16 (2020).

51. Hampshire, A., Highfield, R. R., Parkin, B. L. & Owen, A. M. Fractionating Human Intelligence. Neuron 76, 1225–1237 (2012).

52. Wang, H. et al. A coordinate-based meta-analysis of the n-back working memory paradigm using activation likelihood estimation. Brain Cogn. 132, 1–12 (2019).

53. Haatveit, B. C. et al. The validity of d prime as a working memory index: Results from the “Bergen n-back task”. J. Clin. Exp. Neuropsychol. 32, 871–880 (2010).

54. Corbetta, M. & Shulman, G. L. Spatial neglect and attention networks. Annu. Rev. Neurosci. 34, 569–599 (2011).

55. Corbetta, M. & Shulman, G. L. Control of goal-directed and stimulus-driven attention in the brain. Nat. Rev. Neurosci. 3, 201–215 (2002).

56. Kastner, S. & Ungerleider, L. G. Mechanisms of Visual Attention in the Human Cortex. Annu. Rev. Neurosci. 23, 315–341 (2000).

57. Müri, R. M., Iba-Zizen, M. T., Derosier, C., Cabanis, E. A. & Pierrot-Deseilligny, C. Location of the human posterior eye field with functional magnetic resonance imaging. J. Neurol. Neurosurg. Psychiatry 60, 445–448 (1996).

58. Ptak, R. & Müri, R. M. The parietal cortex and saccade planning: lessons from human lesion studies. Front. Hum. Neurosci. 7, 254 (2013).

59. González-García, C., Flounders, M. W., Chang, R., Baria, A. T. & He, B. J. Content-specific activity in frontoparietal and default-mode networks during prior-guided visual perception. eLife 7, e36068.

60. Hoffman, P., Jefferies, E., Haffey, A., Littlejohns, T. & Lambon Ralph, M. A. Domain-specific control of semantic cognition: A dissociation within patients with semantic working memory deficits. Aphasiology 27, 740–764 (2013).

61. Thompson, H. E. et al. The contribution of executive control to semantic cognition: Convergent evidence from semantic aphasia and executive dysfunction. J. Neuropsychol. 12, 312–340 (2018).

62. Thompson, H. E., Henshall, L. & Jefferies, E. The role of the right hemisphere in semantic control: A case-series comparison of right and left hemisphere stroke. Neuropsychologia 85, 44–61 (2016).

63. Geranmayeh, F., Brownsett, S. L. E. & Wise, R. J. S. Task-induced brain activity in aphasic stroke patients: what is driving recovery? Brain 137, 2632–2648 (2014).

64. Geranmayeh, F., Leech, R. & Wise, R. J. S. Network dysfunction predicts speech production after left hemisphere stroke. Neurology 86, 1296–1305 (2016).

65. Loh, K. K. et al. Cognitive control of orofacial motor and vocal responses in the ventrolateral and dorsomedial human frontal cortex. Proc. Natl. Acad. Sci. 117, 4994–5005 (2020).

66. Seeley, W. W. et al. Dissociable Intrinsic Connectivity Networks for Salience Processing and Executive Control. J. Neurosci. 27, 2349–2356 (2007).

67. Vincent, J. L., Kahn, I., Snyder, A. Z., Raichle, M. E. & Buckner, R. L. Evidence for a Frontoparietal Control System Revealed by Intrinsic Functional Connectivity. J. Neurophysiol. 100, 3328–3342 (2008).

68. Fox, M. D., Corbetta, M., Snyder, A. Z., Vincent, J. L. & Raichle, M. E. Spontaneous neuronal activity distinguishes human dorsal and ventral attention systems. Proc. Natl. Acad. Sci. 103, 10046–10051 (2006).

69. Assem, M., Shashidhara, S., Glasser, M. F. & Duncan, J. Precise Topology of Adjacent Domain-General and Sensory-Biased Regions in the Human Brain. Cereb. Cortex 32, 2521–2537 (2022).

70. Kane, M. J., Conway, A. R. A., Miura, T. K. & Colflesh, G. J. H. Working memory, attention control, and the n-back task: A question of construct validity. J. Exp. Psychol. Learn. Mem. Cogn. 33, 615–622 (2007).

71. Jaeggi, S. M., Buschkuehl, M., Perrig, W. J. & Meier, B. The concurrent validity of the N-back task as a working memory measure. Memory 18, 394–412 (2010).

72. Vatansever, D., Smallwood, J. & Jefferies, E. Varying demands for cognitive control reveals shared neural processes supporting semantic and episodic memory retrieval. Nat. Commun. 12, 2134 (2021).

73. Stampacchia, S. et al. Control the source: Source memory for semantic, spatial and self-related items in patients with LIFG lesions. Cortex 119, 165–183 (2019).

74. Asano, R., Boeckx, C. & Fujita, K. Moving beyond domain-specific versus domain-general options in cognitive neuroscience. Cortex 154, 259–268 (2022).

75. Hodgson, V. J., Lambon Ralph, M. A. & Jackson, R. L. Multiple dimensions underlying the functional organization of the language network. NeuroImage 241, 118444 (2021).

76. Fedorenko, E. & Blank, I. A. Broca’s Area Is Not a Natural Kind. Trends Cogn. Sci. 24, 270–284 (2020).

77. Hodgson, V. J., Lambon Ralph, M. A. & Jackson, R. L. The cross-domain functional organization of posterior lateral temporal cortex: insights from ALE meta-analyses of 7 cognitive domains spanning 12,000 participants. Cereb. Cortex bhac394 (2022) doi:10.1093/cercor/bhac394.

78. Faul, F., Erdfelder, E., Buchner, A. & Lang, A.-G. Statistical power analyses using G*Power 3.1: Tests for correlation and regression analyses. Behav. Res. Methods 41, 1149–1160 (2009).

79. Davis, C. J. N-Watch: A program for deriving neighborhood size and other psycholinguistic statistics. Behav. Res. Methods 37, 65–70 (2005).

80. Halai, A. D., Welbourne, S. R., Embleton, K. & Parkes, L. M. A comparison of dual gradient-echo and spin-echo fMRI of the inferior temporal lobe. Hum. Brain Mapp. 35, 4118–4128 (2014).

81. Visser, M., Jefferies, E. & Lambon Ralph, M. A. Semantic Processing in the Anterior Temporal Lobes: A Meta-analysis of the Functional Neuroimaging Literature. J. Cogn. Neurosci. 22, 1083–1094 (2009).

82. Gorgolewski, K. et al. Nipype: A Flexible, Lightweight and Extensible Neuroimaging Data Processing Framework in Python. Front. Neuroinformatics 5, (2011).

83. Esteban, O. et al. fMRIPrep: a robust preprocessing pipeline for functional MRI. Nat. Methods 16, 111–116 (2019).

84. Tustison, N. J. et al. N4ITK: Improved N3 Bias Correction. IEEE Trans. Med. Imaging 29, 1310–1320 (2010).

85. Avants, B. B., Epstein, C. L., Grossman, M. & Gee, J. C. Symmetric diffeomorphic image registration with cross-correlation: Evaluating automated labeling of elderly and neurodegenerative brain. Med. Image Anal. 12, 26–41 (2008).

86. Zhang, Y., Brady, M. & Smith, S. Segmentation of brain MR images through a hidden Markov random field model and the expectation-maximization algorithm. IEEE Trans. Med. Imaging 20, 45–57 (2001).

87. Jenkinson, M. & Smith, S. A global optimisation method for robust affine registration of brain images. Med. Image Anal. 5, 143–156 (2001).

88. Jenkinson, M., Bannister, P., Brady, M. & Smith, S. Improved Optimization for the Robust and Accurate Linear Registration and Motion Correction of Brain Images. NeuroImage 17, 825–841 (2002).

89. Cox, R. W. & Hyde, J. S. Software tools for analysis and visualization of fMRI data. NMR Biomed. 10, 171–178 (1997).

90. Posse, S. et al. Enhancement of BOLD-contrast sensitivity by single-shot multi-echo functional MR imaging. Magn. Reson. Med. 42, 87–97 (1999).

91. Jaccard, P. The Distribution of the Flora in the Alpine Zone.1. New Phytol. 11, 37–50 (1912).

92. Brett, M., Anton, J.-L., Valabregue, R. & Poline, J.-B. Region of interest analysis using an SPM toolbox. in Region of interest analysis using an SPM toolbox 1 (2002).

