## Supplementary Materials for "Disentangling the neural correlates of semantic and domain-general control: The roles of stimulus domain and task process"

Included in this document:

**Behavioural data:** Supplementary Table 1, Supplementary Figure 1

**Coordinate peak tables:** Supplementary Tables 2-4

**Hard>easy ANOVA analyses:** Supplementary Figures 2-3

**ROI analyses:** Supplementary Tables 5 & 6, Supplementary Figure 4-6

**Task items:** Semantic and non-semantic stimuli

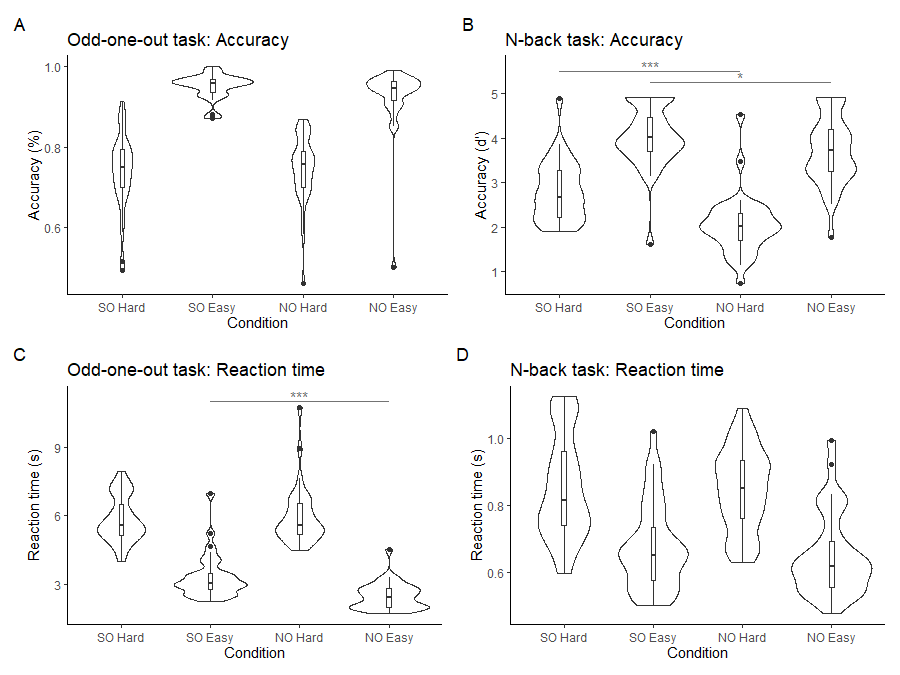
**Behavioural data:** Supplementary Table 1 & Supplementary Figure 1

***Supplementary Figure 1.*** *Violin plots showing behavioural data from the 32 participants across the four task conditions (semantic odd-one-out, SO; non-semantic odd-one-out, NO; semantic* n*-back, SN; non-semantic* n*-back, NN) at both levels of difficulty. A) Performance (accuracy) across the odd-one-out task. B) Performance measured by d-prime (d’) across the* n*-back task, with significant differences between semantic and non-semantic variants at both hard and easy difficulty levels. C) Reaction time across the odd-one-out task, with a significant difference between the semantic and non-semantic variants at the easy difficulty level. D) Reaction time across the* n*-back task.*

| **Supplementary Table 1.** Pairwise significance tests comparing behavioural performance across task-stimulus conditions (32 participants) | |
| --- | --- |
| **Comparison** | **p-value** |
| **Odd-one-out task** | |
| **Accuracy (%)** |  |
| Semantic vs. non-semantic (hard) | 0.801 |
| Semantic vs. non-semantic (easy) | 0.118 |
| Semantic vs. non-semantic (hard-easy) | 0.593 |
| **Reaction time (s)** |  |
| Semantic vs. non-semantic (hard) | 0.422 |
| Semantic vs. non-semantic (easy) | 2.75 x 10-7 *** |
| Semantic vs. non-semantic (hard-easy) | 2.38 x 10-5 *** |
| ***n-*back task** | |
| **Accuracy (d’)** |  |
| Semantic vs. non-semantic (*n*=3) | 8.02 x 10-7 *** |
| Semantic vs. non-semantic (*n*=1) | 0.0112 * |
| Semantic vs. non-semantic (3>1) | 0.0149 * |
| **Reaction time (s)** |  |
| Semantic vs. non-semantic (*n*=3) | 0.938 |
| Semantic vs. non-semantic (*n*=1) | 0.0828 |
| Semantic vs. non-semantic (3>1) | 0.0215 |

**Peak tables:** Supplementary Tables 2-4

| **Supplementary Table 2.** Peak coordinates for activation in the four task-stimulus conditions (hard > rest contrasts) | | | | | | |
| --- | --- | --- | --- | --- | --- | --- |
| **Condition** | **Cluster size** | **T** | **Coordinates** | | | **Brain area** |
|  |  |  | **x** | **y** | **z** |  |
| Semantic odd-one-out | 4191 | 16.2 | -5 | -73 | -27 | Left cerebellum/lingual gyrus |
|  |  | 15.7 | 8 | -71 | -29 |  |
|  |  | 13.3 | 22 | -95 | -5 |  |
|  | 3199 | 14.0 | -21 | -99 | -7 | Inferior occipital/fusiform |
|  |  | 12.5 | -33 | -89 | -15 |  |
|  |  | 12.2 | -45 | -67 | -15 |  |
|  | 3325 | 11.5 | -51 | 30 | 22 | Left IFG/insula |
|  |  | 11.4 | -33 | 24 | -1 |  |
|  |  | 10.6 | -49 | 24 | 28 |  |
|  | 275 | 9.13 | 28 | -65 | 50 | Right angular gyrus |
|  | 567 | 8.52 | -29 | -65 | 48 | Right superior parietal lobe |
|  | 147 | 8.18 | 34 | 26 | -3 | Right insula |
|  | 11 | 6.43 | 22 | -37 | -45 | Right cerebellum |
|  | 48 | 6.36 | -9 | 14 | 50 | Left supplementary motor area |
|  | 41 | 6.34 | -29 | -5 | 58 | Left precentral gyrus |
|  |  | 6.06 | -37 | -3 | 60 |  |
|  | 16 | 6.17 | 44 | 32 | 26 | Right inferior frontal gyrus |
|  | 7 | 6.16 | -47 | 4 | 58 | Left precentral gyrus |
|  | 5 | 5.88 | -11 | -79 | 14 | Left calcarine sulcus |
|  | 2 | 5.79 | -31 | -7 | 66 | Left precentral gyrus |
|  | 1 | 5.75 | -39 | -3 | 56 | Left precentral gyrus |
|  | 1 | 5.74 | -45 | 2 | 60 | Left precentral gyrus |
| Non-semantic odd-one-out | 24948 | 17.2 | -49 | -63 | -7 | Occipital & inferior temporal lobes |
|  |  | 16.8 | 38 | -83 | -11 |  |
|  |  | 16.1 | 14 | -87 | -11 |  |
|  | 242 | 11.9 | -33 | 22 | -3 | Left insula |
|  | 363 | 11.3 | 34 | 26 | -1 | Right insula |
|  | 372 | 9.75 | 50 | 10 | 38 | Right precentral & inferior frontal gyri |
|  |  | 5.94 | 34 | 16 | 32 |  |
|  | 169 | 9.71 | -21 | -39 | -43 | Left cerebellum |
|  | 316 | 9.29 | 44 | 36 | 26 | Right inferior frontal gyrus |
|  | 97 | 9.26 | -29 | -29 | -1 | Left hippocampus |
|  | 598 | 8.96 | -47 | 2 | 32 | Left precentral gyrus |
|  | 107 | 8.76 | 22 | -39 | -45 | Right cerebellum |
|  | 83 | 8.36 | 24 | -27 | -3 | Right hippocampus |
|  | 71 | 7.73 | -47 | 32 | 30 | Left inferior frontal sulcus |
|  | 223 | 7.54 | 28 | -3 | 50 | Right precentral sulcus |
|  |  | 7.22 | 26 | 6 | 52 |  |
|  | 75 | 7.41 | 12 | -7 | 8 | Right supplementary motor area |
|  | 192 | 7.32 | -49 | 52 | 2 | Left frontal pole |
|  |  | 7.19 | -51 | 50 | -7 |  |
|  |  | 6.28 | -53 | 44 | -13 |  |
|  | 124 | 6.76 | -27 | -5 | 56 | Left superior frontal sulcus |
|  | 46 | 6.44 | 32 | 2 | 62 | Right superior frontal sulcus |
|  | 47 | 6.42 | 10 | 20 | 46 | Right cingulum |
|  | 18 | 6.31 | 16 | 14 | 4 | Right caudate |
|  | 6 | 6.12 | -5 | -23 | -11 | Midbrain |
|  | 11 | 6.04 | -13 | 12 | 4 | Left caudate |
|  | 10 | 6.01 | -11 | 14 | 50 | Left supplementary motor area |
|  | 2 | 5.96 | 10 | -17 | -13 | Midbrain |
|  | 2 | 5.93 | -15 | -5 | 10 | Left putamen |
|  | 4 | 5.86 | 20 | -5 | 22 | Right caudate |
|  | 3 | 5.85 | -15 | -55 | -49 | Left cerebellum |
| Semantic n-back | 838 | 14.7 | 36 | 24 | -3 | Right insula |
|  | 637 | 13.3 | -35 | 22 | -3 | Left insula |
|  | 624 | 10.7 | -9 | -75 | -25 | Cerebellum |
|  |  | 9.76 | 10 | -73 | -29 |  |
|  | 824 | 10.5 | -41 | -65 | -29 | Left cerebellum |
|  |  | 8.66 | -35 | -71 | -51 |  |
|  | 360 | 9.87 | 40 | -65 | -31 | Right cerebellum |
|  | 638 | 9.22 | 46 | 34 | 32 | Right middle frontal gyrus |
|  | 1314 | 8.67 | 34 | -69 | 46 | Right parietal lobe/angular gyrus |
|  |  | 7.78 | 38 | -45 | 50 |  |
|  |  | 7.48 | 42 | -53 | 58 |  |
|  | 723 | 7.97 | -45 | 24 | 28 | Left inferior frontal/precentral gyri |
|  |  | 7.49 | -41 | 2 | 38 |  |
|  |  | 7.11 | -39 | 16 | 22 |  |
|  | 521 | 7.94 | 24 | 8 | 62 | Left putamen |
|  |  | 7.79 | 32 | 4 | 62 |  |
|  |  | 7.00 | 40 | 8 | 58 |  |
|  | 154 | 7.92 | -33 | 2 | 56 | Left precentral sulcus |
|  | 472 | 7.89 | -35 | -53 | 42 | Left inferior parietal lobe |
|  |  | 7.65 | -29 | -59 | 42 |  |
|  | 138 | 7.49 | -39 | 54 | 10 | Left anterior middle frontal gyrus |
|  | 70 | 7.39 | 36 | -67 | -53 | Left inferior occipito-temporal lobe |
|  | 414 | 7.38 | 8 | 22 | 42 | Supplementary motor area & anterior cingulate gyrus |
|  |  | 6.77 | -1 | 16 | 54 |  |
|  |  | 6.76 | -11 | 12 | 54 |  |
|  | 183 | 7.02 | 36 | 64 | 4 | Right frontal pole |
|  |  | 6.00 | 30 | 62 | -9 |  |
|  | 28 | 6.47 | 4 | -53 | -15 | Cerebellum |
|  | 17 | 6.32 | 10 | -1 | 2 | Right thalamus |
|  | 13 | 6.14 | -47 | -41 | 46 | Left inferior parietal lobe |
|  | 7 | 5.91 | 34 | 10 | 38 | Right middle frontal gyrus |
|  | 1 | 5.73 | 48 | 14 | 42 | Right inferior frontal sulcus |
|  | 1 | 5.68 | 20 | 14 | 48 | Right superior frontal gyrus |
| Non-semantic n-back | 552 | 12.1 | 32 | 26 | -1 | Right insular/inferior frontal gyrus |
|  |  | 7.35 | 46 | 20 | 4 |  |
|  | 365 | 10.3 | -31 | -61 | -33 | Left cerebellum |
|  |  | 8.01 | -35 | -71 | -51 |  |
|  | 116 | 8.69 | -11 | -77 | -23 | Left cerebellum |
|  | 245 | 8.62 | -31 | 26 | -3 | Left insula |
|  |  | 8.06 | -31 | 20 | 6 |  |
|  | 948 | 8.58 | 32 | -63 | 42 | Right inferior parietal lobe/angular gyrus |
|  |  | 8.08 | 38 | -47 | 48 |  |
|  |  | 7.13 | 48 | -37 | 52 |  |
|  | 223 | 7.97 | 44 | 34 | 34 | Right middle frontal gyrus |
|  | 62 | 7.57 | -11 | 16 | 48 | Left supplementary motor area |
|  | 276 | 7.15 | -29 | -55 | 44 | Left inferior parietal lobe |
|  |  | 5.89 | -29 | -63 | 38 |  |
|  | 37 | 7.02 | 10 | 22 | 42 | Right anterior cingulate gyrus |
|  | 34 | 6.78 | -29 | -3 | 58 | Left precentral sulcus |
|  | 20 | 6.63 | 40 | 44 | 22 | Right middle frontal gyrus |
|  | 27 | 6.41 | 28 | -61 | -31 | Right cerebellum |
|  | 20 | 6.34 | -13 | -101 | 8 | Left middle occipital gyrus |
|  | 13 | 6.27 | -45 | 26 | 30 | Left inferior frontal sulcus |
|  | 35 | 6.12 | 16 | -97 | 10 | Right inferior occipital lobe/cuneus |
|  | 9 | 6.07 | -33 | 20 | 26 | Left inferior frontal lobe |
|  | 39 | 6.06 | -45 | -67 | -7 | Left inferior occipital gyrus |
|  | 6 | 6.04 | 34 | -69 | -53 | Right cerebellum |
|  | 9 | 5.83 | 28 | 8 | 52 | Right middle frontal gyrus |
|  | 10 | 5.80 | -47 | -3 | 34 | Left precentral gyrus |
|  | 4 | 5.80 | 30 | 2 | 62 | Right superior frontal sulcus |
|  | 2 | 5.74 | 8 | 16 | 50 | Right supplementary motor area |
|  | 3 | 5.69 | 10 | -73 | -23 | Right cerebellum |

| **Supplementary Table 3**. Peak coordinates for activation in the ANOVA (hard > rest contrasts) | | | | | |
| --- | --- | --- | --- | --- | --- |
|  | **Cluster size** | **F statistic** | **Coordinates** | | |
|  |  |  | **x** | **y** | **z** |
| Main effect of task | 15107 | 225 | 24 | -95 | -7 |
|  |  | 220 | 34 | -85 | -7 |
|  |  | 209 | -27 | -93 | -9 |
|  | 3867 | 106 | 58 | -51 | 46 |
|  |  | 105 | 54 | -21 | -9 |
|  |  | 94.1 | 54 | -47 | 32 |
|  | 790 | 92.2 | -3 | -73 | -27 |
|  |  | 78.0 | -1 | -57 | -37 |
|  |  | 37.3 | 14 | -75 | -45 |
|  | 186 | 86.7 | 22 | -39 | -45 |
|  | 193 | 82.8 | -21 | -39 | -45 |
|  | 264 | 81.7 | -21 | -31 | -3 |
|  | 1726 | 68.7 | -53 | 32 | 18 |
|  |  | 61.3 | -51 | 38 | 12 |
|  |  | 59.8 | -43 | 36 | -13 |
|  | 1155 | 66.1 | 32 | 34 | 42 |
|  |  | 48.5 | 22 | 58 | 24 |
|  |  | 37.9 | 24 | 64 | 2 |
|  | 597 | 59.8 | 8 | 40 | 22 |
|  |  | 44.7 | 12 | 42 | -1 |
|  | 670 | 57.9 | 4 | -25 | 32 |
|  |  | 48.0 | 12 | -65 | 36 |
|  |  | 43.0 | 8 | -43 | 40 |
|  | 109 | 57.4 | 24 | -27 | -3 |
|  | 869 | 50.1 | -55 | -27 | 42 |
|  |  | 38.8 | -43 | -25 | 62 |
|  |  | 38.6 | -43 | -35 | 52 |
|  | 114 | 41.2 | 34 | 16 | -9 |
|  |  | 31.2 | 46 | 18 | 2 |
|  | 165 | 38.4 | -45 | -59 | -43 |
|  | 93 | 37.6 | 46 | -25 | 36 |
|  | 21 | 37.4 | -33 | 16 | -11 |
|  | 105 | 37.2 | 16 | 12 | 64 |
|  | 148 | 36.1 | -53 | -57 | 40 |
|  |  | 27.5 | -61 | -49 | 46 |
|  | 68 | 33.8 | -57 | -51 | 18 |
|  | 40 | 32.0 | -29 | 36 | 40 |
|  | 23 | 31.2 | -31 | 52 | 24 |
|  | 33 | 28.9 | 34 | -73 | -41 |
|  | 7 | 27.3 | 50 | 10 | 32 |
|  | 3 | 27.0 | 20 | 20 | 58 |
|  | 6 | 26.2 | -11 | -69 | 34 |
|  | 4 | 25.7 | 26 | 20 | 58 |
|  | 4 | 25.6 | -9 | -45 | 46 |
|  | 4 | 25.5 | 8 | -75 | -13 |
| Main effect of stimulus | 6669 | 120 | 34 | -85 | 10 |
|  |  | 112 | 44 | -61 | -11 |
|  |  | 106 | 46 | -53 | -7 |
|  | 3248 | 83.2 | -29 | -57 | -11 |
|  |  | 75.6 | -29 | -91 | 20 |
|  |  | 69.9 | -33 | -87 | 10 |
|  | 658 | 64.1 | -59 | -43 | 6 |
|  | 377 | 43.4 | -49 | 32 | 2 |
|  |  | 28.6 | -51 | 28 | 16 |
|  | 27 | 33.6 | -17 | -47 | -51 |
|  | 21 | 29.9 | -49 | 18 | 24 |
|  | 7 | 27.0 | 14 | -83 | -35 |
|  | 1 | 25.6 | 20 | -31 | 2 |
| Interaction task x stimulus | 671 | 46.5 | 30 | -73 | 34 |
|  |  | 38.9 | 36 | -83 | 24 |
|  |  | 31.2 | 34 | -85 | 8 |
|  | 545 | 42.5 | 48 | -53 | -7 |
|  |  | 38.0 | 44 | -61 | -11 |
|  |  | 33.5 | 54 | -63 | -3 |
|  | 169 | 38.5 | 40 | -41 | 50 |
|  |  | 29.9 | 42 | -45 | 58 |
|  | 60 | 34.1 | -19 | -75 | -49 |
|  |  | 30.9 | -29 | -73 | -51 |
|  | 72 | 30.7 | -33 | -83 | 24 |
|  |  | 26.8 | -25 | -81 | 36 |
|  | 13 | 27.5 | 12 | -71 | 56 |
|  | 17 | 27.3 | -13 | -67 | 54 |
|  | 3 | 26.1 | 10 | -89 | -9 |
|  | 1 | 25.4 | 58 | -27 | 50 |
|  | 1 | 25.3 | 24 | 10 | 62 |
|  | 2 | 25.0 | 50 | 10 | 36 |
|  | 1 | 24.8 | -9 | -53 | 14 |

| **Supplementary Table 4.** Peak coordinates for post-hoc comparison tests in the ANOVA (hard > rest contrasts) | | | | | |
| --- | --- | --- | --- | --- | --- |
| **Contrast** | **Cluster size** | **T statistic** | **Coordinates** | | |
|  |  |  | **x** | **y** | **z** |
| Semantic > non-semantic stimuli | 1434 | 7.76 | -51 | 20 | 24 |
|  |  | 7.12 | -47 | 32 | 4 |
|  |  | 6.93 | -55 | 24 | 2 |
|  | 962 | 7.65 | -59 | -41 | 6 |
|  |  | 7.05 | -59 | -53 | 6 |
|  |  | 7.01 | -59 | -31 | 2 |
|  | 142 | 6.75 | 14 | -83 | -33 |
|  | 61 | 6.17 | -45 | -51 | -19 |
| Non-semantic > semantic stimuli | 7710 | 10.7 | 36 | -83 | 10 |
|  |  | 10.1 | 30 | -45 | -13 |
|  |  | 10.0 | 44 | -59 | -9 |
|  | 3973 | 10.4 | -29 | -53 | -13 |
|  |  | 8.78 | -31 | -91 | 14 |
|  |  | 8.36 | -23 | -97 | 16 |
|  | 28 | 5.78 | -17 | -45 | -51 |
|  | 46 | 5.41 | -17 | -67 | 44 |
|  | 7 | 5.35 | -25 | -31 | -3 |
| Odd-one-out > n-back task | 19769 | 18.5 | -27 | -95 | -9 |
|  |  | 17.9 | 22 | -97 | -5 |
|  |  | 13.4 | -39 | -81 | -13 |
|  |  | 12.6 | -51 | -55 | -13 |
|  | 2688 | 11.4 | -51 | 40 | 10 |
|  |  | 8.91 | -51 | 8 | 30 |
|  |  | 8.57 | -49 | 40 | -13 |
|  |  | 7.01 | -57 | 16 | 26 |
|  | 343 | 9.78 | -21 | -31 | -1 |
|  | 211 | 9.44 | 22 | -39 | -45 |
|  | 202 | 9.03 | -23 | -37 | -45 |
|  | 114 | 7.62 | 24 | -27 | -3 |
|  | 421 | 6.90 | 42 | -27 | 40 |
|  | 163 | 6.81 | 44 | 34 | 18 |
|  | 56 | 5.97 | 34 | -71 | -43 |
|  | 126 | 5.91 | -7 | 54 | -19 |
|  |  | 5.84 | -7 | 42 | -21 |
|  | 22 | 5.77 | 30 | 34 | -19 |
|  | 10 | 5.60 | -39 | -5 | 16 |
|  | 17 | 5.42 | 52 | 10 | 32 |
|  | 12 | 5.35 | -23 | -11 | 48 |
|  | 3 | 5.21 | 26 | -59 | -55 |
|  | 3 | 5.10 | 10 | -73 | -11 |
|  | 1 | 5.09 | -39 | -13 | 56 |
| n-back > odd-one-out task | 4419 | 10.4 | 56 | -51 | 44 |
|  |  | 9.92 | 52 | -49 | 36 |
|  |  | 9.85 | 62 | -23 | -9 |
|  |  | 8.68 | 60 | -45 | 12 |
|  | 3744 | 8.84 | 10 | 38 | 24 |
|  |  | 8.05 | 10 | 50 | -1 |
|  |  | 7.95 | 24 | 60 | 22 |
|  |  | 7.78 | 26 | 64 | 2 |
|  | 1563 | 8.63 | 12 | -63 | 38 |
|  |  | 8.18 | 6 | -25 | 32 |
|  |  | 7.65 | 8 | -41 | 40 |
|  | 368 | 8.02 | 34 | 18 | -9 |
|  |  | 6.88 | 44 | 18 | 4 |
|  | 365 | 7.56 | -43 | -57 | -41 |
|  | 277 | 7.28 | -31 | 52 | 22 |
|  |  | 5.70 | -29 | 54 | 8 |
|  | 569 | 7.22 | -55 | -51 | 18 |
|  |  | 6.34 | -47 | -55 | 44 |
|  |  | 6.19 | -55 | -51 | 40 |
|  | 140 | 6.87 | -9 | -67 | 38 |
|  |  | 5.30 | -11 | -57 | 42 |
|  | 39 | 6.59 | -33 | 16 | -11 |
|  | 99 | 6.16 | 14 | 14 | 64 |
|  | 219 | 6.07 | -31 | 36 | 38 |
|  | 53 | 5.89 | -53 | -29 | -9 |
|  | 8 | 5.68 | -33 | -55 | 6 |
|  | 4 | 5.38 | -7 | -55 | -55 |
|  | 4 | 5.18 | -7 | -61 | 50 |
| Semantic odd-one-out > non-semantic odd-one-out | 941 | 9.83 | -41 | 32 | -15 |
|  |  | 8.08 | -23 | 30 | -13 |
|  |  | 7.67 | -47 | 32 | 6 |
|  |  | 7.09 | -57 | 18 | 26 |
|  | 254 | 7.09 | -63 | -41 | 6 |
|  |  | 6.72 | -63 | -49 | 2 |
|  | 40 | 6.88 | -7 | -57 | 16 |
|  | 49 | 6.83 | 14 | -83 | -35 |
|  | 17 | 6.39 | -45 | -51 | -21 |
|  | 25 | 6.08 | -51 | -17 | -15 |
|  |  | 5.86 | -57 | -11 | -15 |
|  | 6 | 6.08 | 22 | -43 | 16 |
|  | 3 | 5.88 | -9 | 30 | -17 |
|  | 1 | 5.72 | -39 | 22 | -27 |
|  | 1 | 5.71 | -29 | -13 | -17 |
| Non-semantic odd-one-out > semantic odd-one-out | 12527 | 15.9 | 48 | -55 | -7 |
|  |  | 14.7 | 30 | -85 | 10 |
|  |  | 13.1 | 38 | -83 | 20 |
|  |  | 12.3 | 36 | -77 | 28 |
|  | 885 | 11.8 | -29 | -55 | -11 |
|  |  | 7.45 | -19 | -81 | -17 |
|  |  | 6.42 | -21 | -77 | -5 |
|  | 127 | 7.80 | -21 | -73 | -47 |
|  |  | 6.13 | -9 | -77 | -41 |
|  | 57 | 6.99 | -15 | -53 | -51 |
|  | 39 | 6.62 | 50 | 8 | 32 |
|  | 22 | 6.30 | -33 | -53 | 58 |
|  | 19 | 6.22 | 58 | -25 | 46 |
|  | 11 | 6.19 | 20 | -31 | 6 |
|  | 2 | 6.02 | 16 | -25 | 14 |
| Semantic n-back > non-semantic n-back | 1 | 5.77 | -59 | -29 | 2 |
|  | 1 | 5.62 | -19 | 4 | 58 |
| Non-semantic n-back > semantic n-back | 88 | 6.59 | 40 | -81 | 10 |
|  | 7 | 6.09 | 38 | -45 | -9 |
|  | 3 | 5.69 | -27 | -51 | -13 |
| Semantic odd-one-out > semantic n-back | 1169 | 12.6 | 24 | -97 | -7 |
|  |  | 7.37 | 46 | -79 | -7 |
|  |  | 6.88 | 36 | -85 | 10 |
|  | 1739 | 12.6 | -27 | -95 | -11 |
|  |  | 9.98 | -51 | -55 | -13 |
|  |  | 9.09 | -39 | -79 | -15 |
|  | 1511 | 11.7 | -49 | 38 | 8 |
|  |  | 9.09 | -41 | 32 | -13 |
|  |  | 7.48 | -27 | 30 | -15 |
|  |  | 6.67 | -57 | 20 | 20 |
|  | 120 | 8.03 | -33 | -33 | -21 |
|  | 69 | 7.00 | -1 | -73 | -29 |
|  |  | 6.07 | 10 | -81 | -33 |
|  | 19 | 6.85 | 22 | -39 | -45 |
|  | 73 | 6.50 | -13 | -79 | 14 |
|  |  | 5.95 | -21 | -69 | 8 |
|  | 23 | 6.39 | -15 | -53 | 4 |
|  | 14 | 6.27 | -21 | -31 | -1 |
|  | 21 | 6.18 | -27 | -73 | 40 |
|  | 26 | 6.17 | -29 | -79 | 18 |
|  | 15 | 6.13 | -5 | 36 | -21 |
|  | 26 | 6.06 | 16 | -63 | 8 |
|  | 2 | 5.74 | 16 | -75 | 16 |
| Semantic n-back > semantic odd-one-out | 2265 | 9.70 | 52 | -49 | 48 |
|  |  | 8.76 | 42 | -77 | 42 |
|  |  | 8.73 | 44 | -69 | 44 |
|  |  | 6.86 | 62 | -45 | 18 |
|  | 115 | 8.69 | 12 | 34 | 28 |
|  | 1277 | 8.55 | 40 | 46 | 24 |
|  |  | 8.04 | 32 | 32 | 46 |
|  |  | 7.97 | 40 | 40 | 34 |
|  |  | 7.70 | 30 | 64 | 2 |
|  | 193 | 8.41 | 8 | -67 | 42 |
|  | 295 | 8.01 | -43 | -63 | -43 |
|  |  | 6.68 | -39 | -45 | -41 |
|  |  | 6.61 | -37 | -63 | -33 |
|  | 291 | 7.21 | 24 | 16 | 48 |
|  |  | 7.07 | 22 | 10 | 66 |
|  |  | 6.41 | 22 | 14 | 56 |
|  | 109 | 7.15 | 34 | 18 | -9 |
|  |  | 6.52 | 44 | 20 | 4 |
|  | 220 | 7.09 | 58 | -21 | -9 |
|  | 35 | 6.99 | -7 | -63 | 52 |
|  | 154 | 6.89 | -51 | -53 | 38 |
|  |  | 6.09 | -47 | -53 | 46 |
|  | 39 | 6.86 | -31 | 52 | 22 |
|  | 111 | 6.47 | 8 | -47 | 46 |
|  | 7 | 6.41 | -7 | -57 | -55 |
|  | 18 | 6.08 | -33 | 32 | 42 |
|  | 10 | 6.06 | -9 | -69 | 40 |
|  | 3 | 5.95 | 6 | -25 | 30 |
|  | 5 | 5.91 | -33 | 18 | -11 |
|  | 3 | 5.81 | -51 | -53 | 16 |
|  | 3 | 5.77 | 10 | 50 | -1 |
| Non-semantic odd-one-out > non-semantic n-back | 19620 | 15.4 | 48 | -55 | -7 |
|  |  | 15.0 | -27 | -95 | -7 |
|  |  | 14.1 | 40 | -85 | 18 |
|  |  | 14.0 | -35 | -91 | -5 |
|  | 154 | 8.68 | -1 | -59 | -37 |
|  | 260 | 8.42 | -51 | 6 | 30 |
|  | 169 | 8.41 | -23 | -41 | -45 |
|  | 124 | 8.16 | -27 | -29 | -3 |
|  | 169 | 7.95 | 52 | 10 | 34 |
|  | 71 | 7.64 | 24 | -27 | -3 |
|  | 141 | 7.54 | 22 | -41 | -47 |
|  | 85 | 7.12 | 26 | -5 | 46 |
|  | 47 | 6.72 | -53 | 42 | 12 |
|  | 47 | 6.53 | 44 | 36 | 18 |
|  | 29 | 6.34 | 14 | -75 | -47 |
|  | 4 | 5.84 | -51 | 42 | -15 |
| Non-semantic n-back > non-semantic odd-one-out | 363 | 9.30 | 68 | -21 | -11 |
|  | 177 | 7.37 | 60 | -53 | 46 |
|  |  | 6.54 | 54 | -51 | 34 |
|  | 297 | 7.24 | 12 | 44 | 14 |
|  |  | 6.38 | 12 | 58 | -1 |
|  |  | 6.10 | 8 | 44 | -3 |
|  | 33 | 6.69 | 6 | -25 | 36 |
|  | 45 | 6.19 | 50 | -43 | 6 |
|  |  | 6.04 | 56 | -47 | 14 |
|  | 12 | 6.18 | 22 | 56 | 20 |
|  | 10 | 5.93 | 66 | -47 | 32 |
|  | 1 | 5.69 | 6 | -41 | 30 |

**Additional ANOVA analyses:** Supplementary Figures 2-3

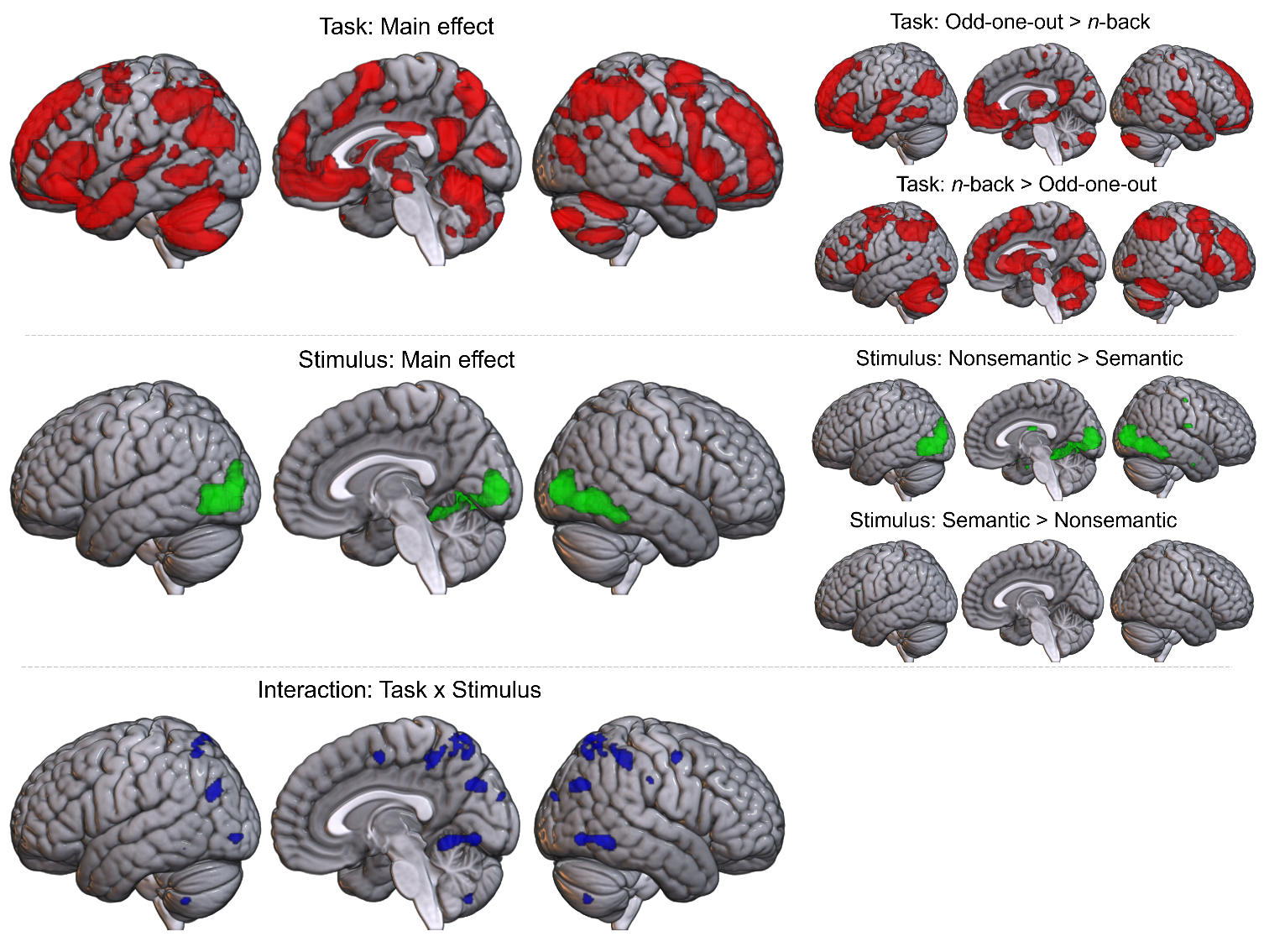

***Supplementary Figure 2.*** *Results of a two-way ANOVA at the second level, constructed using the hard > easy first level contrasts. Voxel-level threshold p<0.05 with FWE correction. Top box: main effect of task (odd-one-out vs.* n*-back) and associated post-hoc tests. Middle box: main effect of stimulus (semantic vs non-semantic) and associated post-hoc tests. Lower box: interaction of task and stimulus. While the results are similar to the hard>rest analyses, the large difficulty differences between the two tasks dominates the task effect which also reduces the stimuli effect.*

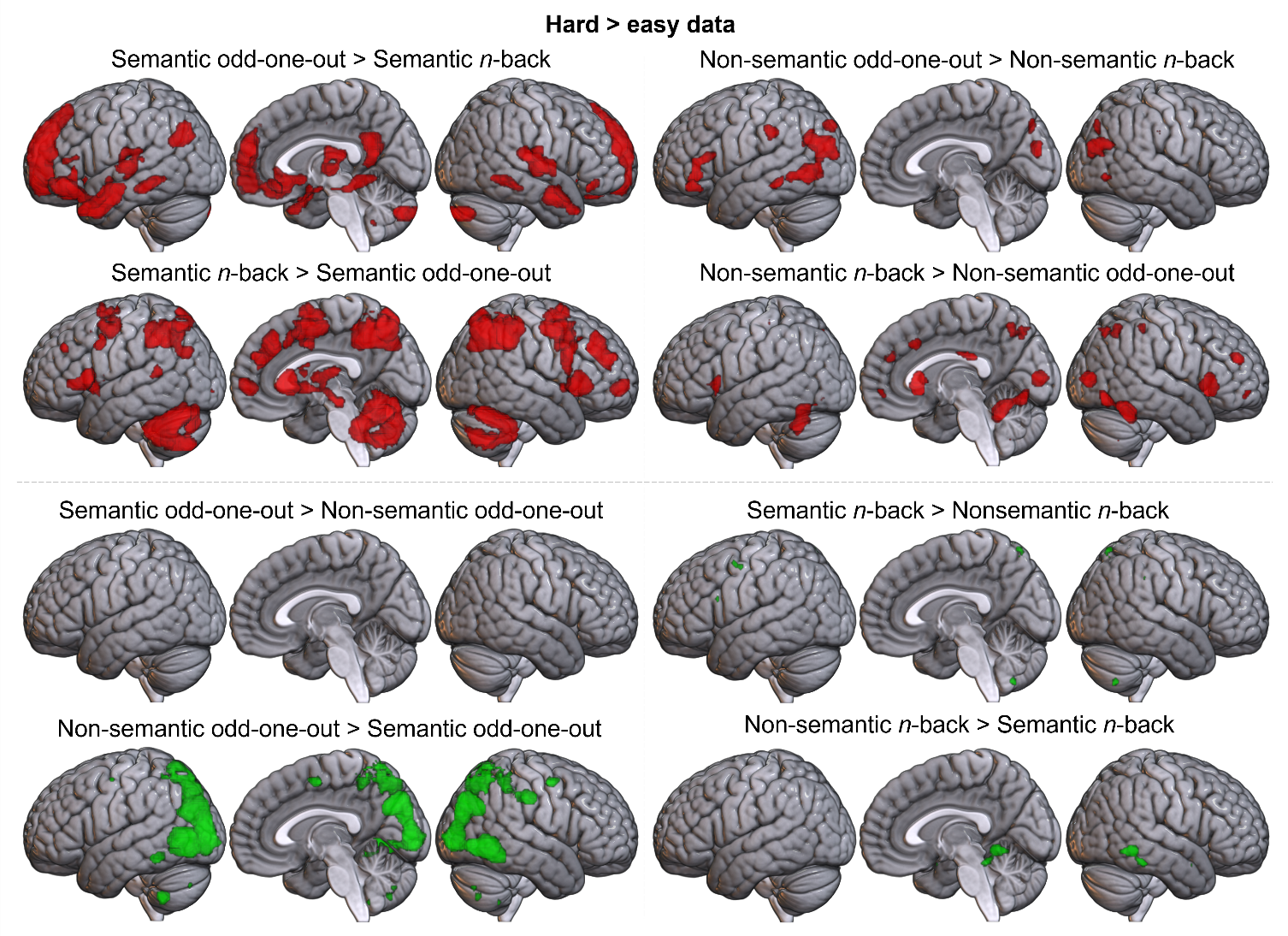

***Supplementary Figure 3.*** *Pairwise comparisons between conditions for the hard>easy contrast. Voxel-level threshold p<0.05 with FWE correction. Top left quadrant: semantic odd-one-out vs.* n*-back task. Top right quadrant: non-semantic odd-one-out vs.* n*-back task. Bottom left quadrant: semantic versus non-semantic odd-one-out task. Bottom right quadrant: semantic versus non-semantic* n*-back task. Comparisons assessing the impact of stimulus domain are shown in green, while comparisons assessing task process effects are displayed in red.*

**ROI analyses:** Supplementary Tables 5-6, Supplementary Figures 4-6

| **Supplementary Table 5.** Results of within-ROI factorial ANOVAs | | | | | | |
| --- | --- | --- | --- | --- | --- | --- |
| **ROI** | **Interaction effect** | | **Stimulus main effect** | | **Task main effect** | |
|  | **F** | **p** | **F** | **p** | **F** | **p** |
| *SCN ROIs* | | | | | | |
| Left IFG (tri.) | 4.19 | 0.0427 * |  | |  | |
| Left IFG (orb.) | 0.434 | 0.511 | 27.5 | 6.52 x 10^-7^ *** | 3.66 | 0.0581 |
| Right IFG (tri.) | 2.72 | 0.102 | 0.0499 | 0.824 | 1.49 | 0.225 |
| Right IFG (orb.) | 4.50 | 0.0359 * |  | |  | |
| pMTG/STS | 8.25 | 0.00480 ** |  | |  | |
| pITG | 1.90 | 0.171 | 17.1 | 6.58 x 10^-5^ *** | 55.6 | 1.33 x 10^-11^ *** |
| dmPFC | 0.400 | 0.528 | 2.80 | 0.0965 | 0.749 | 0.389 |
| *MDN ROIs* | | | | | | |
| Left SFG | 9.11 | 0.00302 ** |  | |  | |
| Right SFG | 13.2 | 0.000406 *** |  | |  | |
| Left PCG | 3.94 | 0.0493 * |  | |  | |
| Right PCG | 14.1 | 0.000271 *** |  | |  | |
| Left IPL | 6.77 | 0.0104 * |  | |  | |
| Right IPL | 15.9 | 0.000112 *** |  | |  | |
| Left IOC | 7.73 | 0.00629 ** |  | |  | |
| Right IOC | 19.0 | 2.73 x 10^-5^ *** |  | |  | |
| Left insula | 3.87 | 0.514 | 1.78 | 0.185 | 0.561 | 0.455 |
| Right insula | 5.93 | 0.0163 * |  |  |  |  |
| SMA | 0.709 | 0.401 | 2.33 | 0.130 | 0.124 | 0.725 |
| Left aMFG | 1.09 | 0.297 | 0.382 | 0.537 | 7.19 | 0.00833 ** |
| Right aMFG | 5.84 | 0.0171 * |  |  |  |  |
| Left mid MFG | 0.779 | 0.379 | 7.34 | 0.00768 ** | 5.66 | 0.0189 * |
| Right mid MFG | 6.93 | 0.00953 ** |  |  |  |  |
| *Main effects are not reported in the context of a significant interaction. Instead, the results of multiple-comparison post-hoc pairwise tests (Tukey’s HSD) are reported in the main text; see Figures 6 & 7 and accompanying discussion.* | | | | | | |

| **Supplementary Table 6.** Coordinates used to create 8mm regions of interest (ROIs) | | | | |
| --- | --- | --- | --- | --- |
| ***Network*** | ***Region*** | ***Coordinates*** | | |
|  |  | ***x*** | ***y*** | ***z*** |
| SCN | Left inferior frontal gyrus (IFG) (pars triangularis) | -48 | 22 | 20 |
|  | Right inferior frontal gyrus (pars triangularis) | 50 | 24 | 26 |
|  | Left inferior frontal gyrus (pars orbitalis) | -46 | 24 | -2 |
|  | Right inferior frontal gyrus (pars orbitalis) | 32 | 24 | -6 |
|  | Left posterior middle temporal gyrus/superior temporal sulcus (pMTG/STS) | -54 | -42 | 4 |
|  | Left posterior inferior temporal gyrus (pITG) | -46 | -48 | -16 |
|  | Dorsomedial prefrontal cortex (dmPFC) | -2 | 20 | 52 |
| MDN | Left superior frontal gyrus (SFG) | -26 | -4 | 56 |
|  | Right superior frontal gyrus | 26 | -4 | 56 |
|  | Left precentral gyrus (PCG) | -46 | 4 | 32 |
|  | Right precentral gyrus | 46 | 4 | 32 |
|  | Left inferior parietal lobule (IPL) | -36 | -46 | 42 |
|  | Right inferior parietal lobule | 36 | -46 | 42 |
|  | Left inferior occipital cortex (IOC) | -34 | -78 | -6 |
|  | Right inferior occipital cortex | 34 | -78 | -6 |
|  | Left insula | -30 | 22 | 4 |
|  | Right insula | 30 | 22 | 4 |
|  | Supplementary motor area (SMA) | 0 | 12 | 50 |
|  | Left anterior middle frontal gyrus (MFG) | -37 | 49 | 18 |
|  | Right anterior middle frontal gyrus | 37 | 49 | 18 |
|  | Left mid middle frontal gyrus | -44 | 32 | 28 |
|  | Right mid middle frontal gyrus | 44 | 32 | 28 |

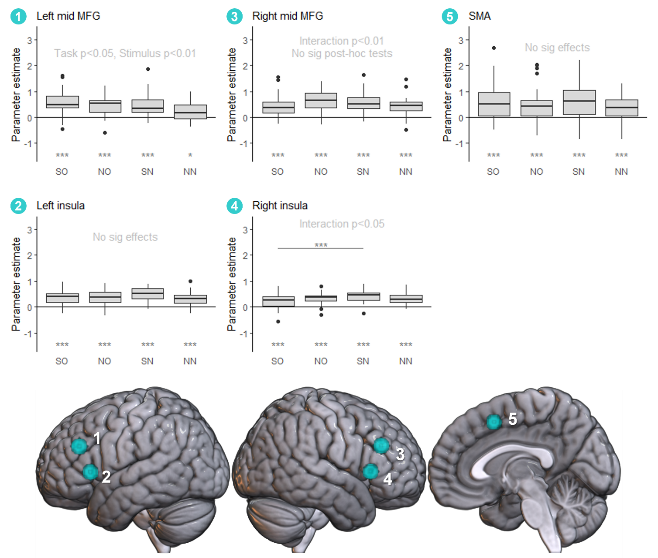

***Supplementary Figure 4.*** *Functional profiles of the additional MDN ROIs. Each ROI is an 8mm sphere around a peak taken from the MDN mask by Fedorenko et al. 2013, with locations shown in the bottom panel. These are not shown in the main text as they overlap SCN ROIs, leading to repetition. Activity for each condition is shown for each ROI (semantic odd-one-out, SO; non-semantic odd-one-out, NO; semantic* n*-back, SN; non-semantic* n*-back, SN). Results of two-way ANOVAs (task process x stimulus modality) for each ROI are shown above where significant, along with the results of multiple comparison corrected post-hoc tests (Tukey’s HSD) between conditions where a significant interaction was observed. Results of multiple comparison corrected one-sample t-tests, to determine whether activation is significantly different from rest in each ROI, are shown with asterisks at the bottom of each graph.*

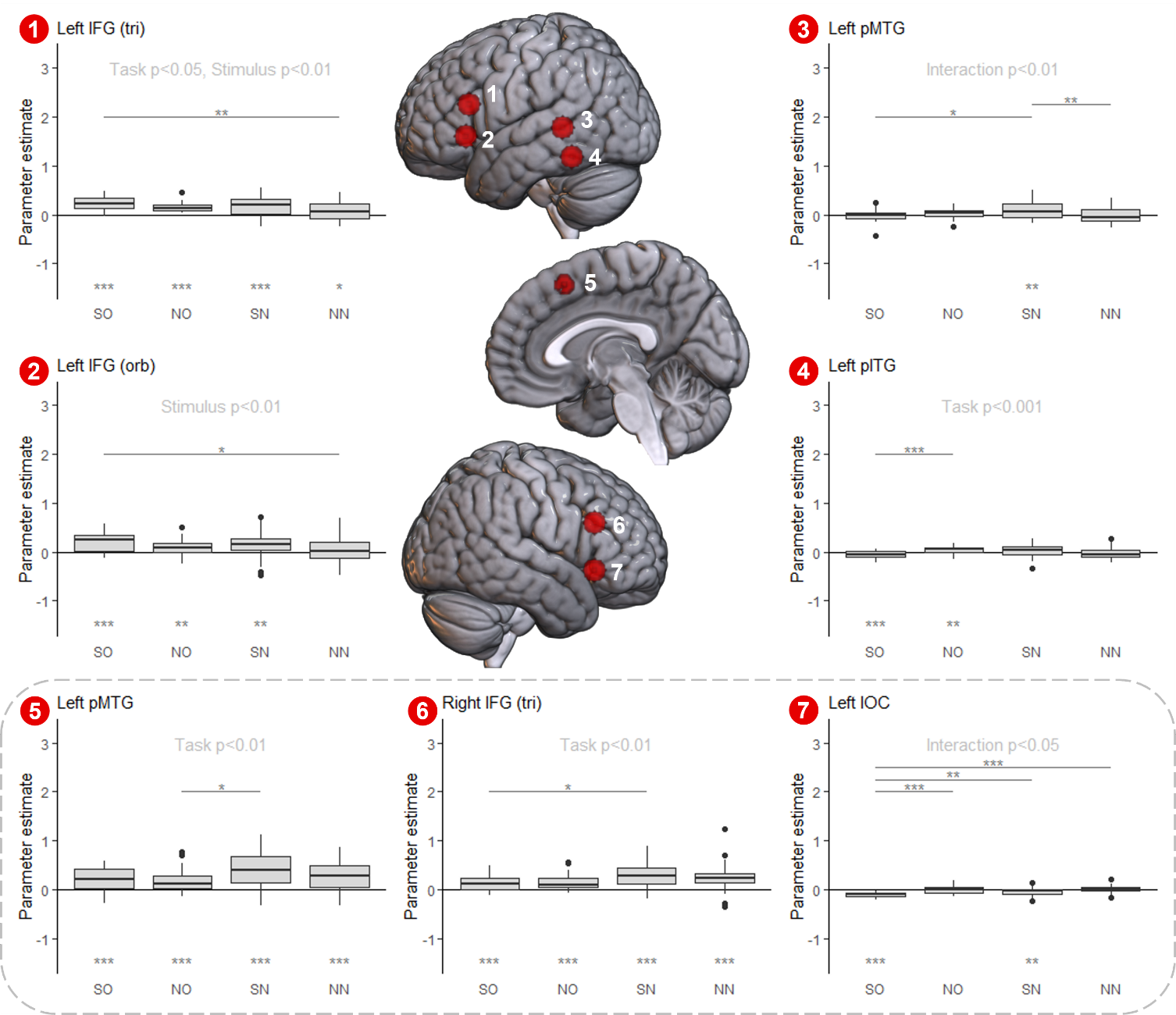

***Supplementary Figure 5.*** *Functional profile of the ROIs derived from the SCN mask, based on the hard>easy contrasts. The location of each ROI is indicated in the centre panel. Activity for each condition is shown for each ROI (semantic odd-one-out, SO; non-semantic odd-one-out, NO; semantic* n*-back, SN; non-semantic* n*-back, SN). Results of a two-way ANOVA (task process x stimulus modality) for each ROI are shown above where significant, along with the results of multiple comparison corrected post-hoc tests (Tukey’s HSD) between conditions where a significant interaction was observed. Significant multiple comparison corrected one-sample t-tests, showing activation is significantly different from rest are shown below with asterisks. Significance levels are indicated with asterisks; *** p<0.001, ** p<0.01, * p<0.05.*

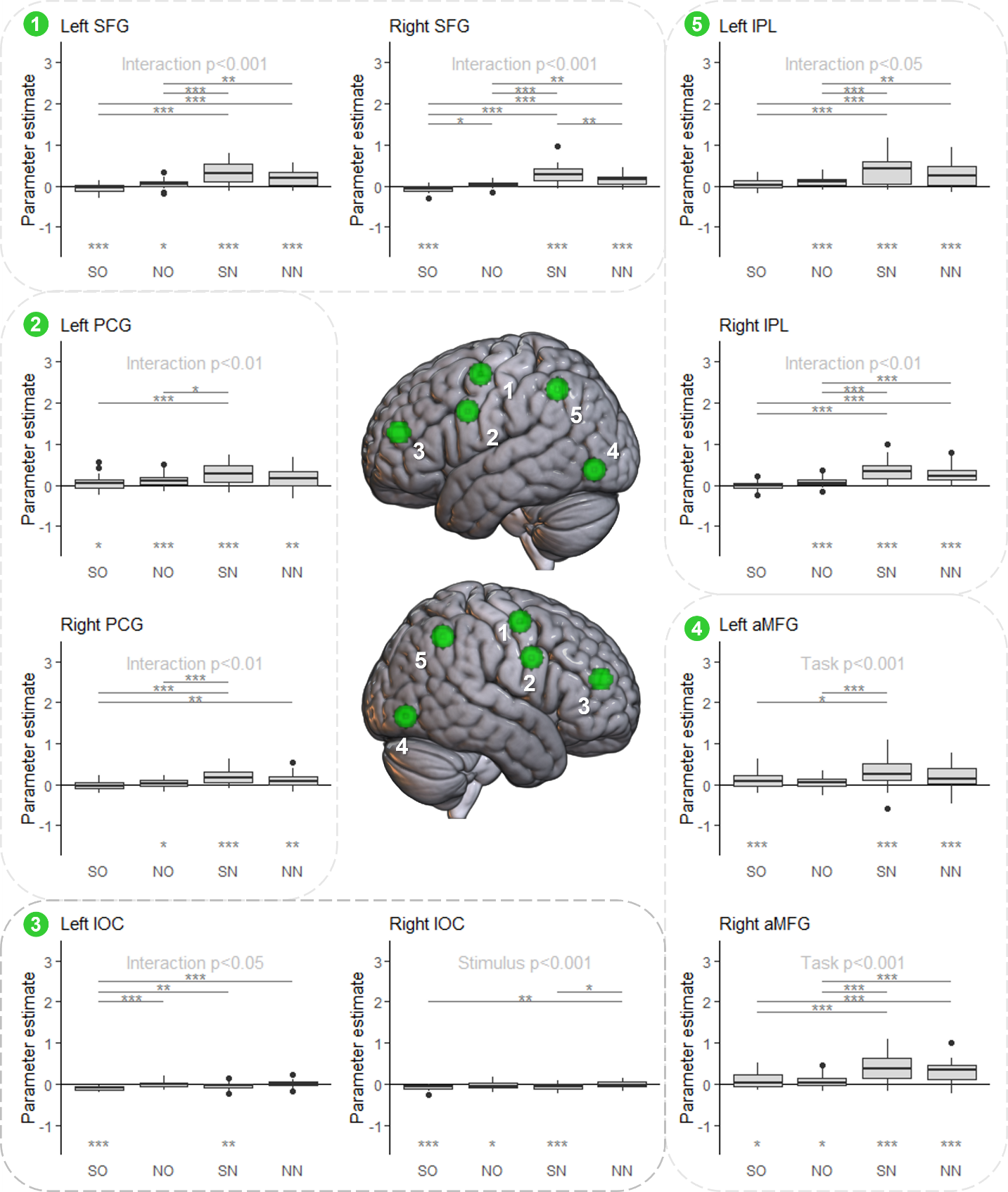

***Supplementary Figure 6.*** *Functional profile of the ROIs derived from the MDN mask, based on the hard>easy contrasts. The location of each ROI is indicated in the centre panel. Activity for each condition is shown for each ROI (semantic odd-one-out, SO; non-semantic odd-one-out, NO; semantic* n*-back, SN; non-semantic* n*-back, SN). Results of two-way ANOVAs (task process x stimulus modality) for each ROI are shown above where significant, along with the results of multiple comparison corrected post-hoc tests (Tukey’s HSD) between conditions where a significant interaction was observed. Significant multiple comparison corrected one-sample t-tests, showing activation is significantly different from rest are shown below with asterisks. Significance levels are indicated with asterisks; *** p<0.001, ** p<0.01, * p<0.05.*

**Task items**

| **Hard semantic odd-one-out trials** | | | | |
| --- | --- | --- | --- | --- |
| ***Item 1*** | ***Item 2*** | ***Item 3*** | ***Item 4*** | ***Correct Answer*** |
| violin | harp | piano | trumpet | 4 |
| coyote | turtle | rabbit | shark | 4 |
| beer | spirit | ghost | wine | 3 |
| violet | army | navy | orange | 2 |
| sink | sail | kettle | knife | 2 |
| foot | metre | elbow | yard | 3 |
| perch | pigeon | sole | tuna | 2 |
| ribbon | bow | quiver | target | 1 |
| spring | fall | drop | trip | 1 |
| ship | craft | art | dinghy | 3 |
| screen | rat | circuit | mouse | 2 |
| mash | fry | bake | boat | 4 |
| lobster | cloud | tomato | ruby | 2 |
| captain | chest | rum | slip | 4 |
| lizard | lion | clown | magician | 1 |
| arrow | needle | thread | thorn | 3 |
| bell | dish | pedal | spoke | 2 |
| chapel | coffin | ring | veil | 2 |
| dwarf | apple | mirror | autumn | 4 |
| coin | hammer | shot | javelin | 1 |
| collar | shirt | bark | paw | 2 |
| emerald | grass | frog | tiger | 4 |
| brush | tinsel | star | diamond | 1 |
| cotton | fire | desert | chilli | 1 |
| club | spade | glove | heart | 3 |
| banana | jeans | whale | sky | 1 |
| bus | car | train | plane | 2 |
| slipper | pumpkin | blanket | midnight | 3 |
| daffodil | lemon | mole | cheese | 3 |
| fox | carrot | basketball | guitar | 4 |
| squash | beetle | cricket | rugby | 2 |
| eagle | bunker | hawk | penguin | 2 |
| psalm | font | capital | aisle | 3 |
| ice | wax | butter | pot | 4 |
| jewel | yacht | mansion | mallet | 4 |
| pillow | petal | walnut | velvet | 3 |
| shoe | cart | dog | race | 3 |
| cow | orchard | tractor | forest | 4 |
| screw | key | mug | handle | 3 |
| parrot | zebra | panther | giraffe | 1 |
| towel | shell | paint | palm | 3 |
| pound | dollar | kilo | ounce | 2 |
| dice | sugar | box | jungle | 4 |
| beaker | swing | bench | picnic | 1 |
| boot | saddle | axe | saloon | 3 |
| panda | domino | wasp | football | 3 |
| cracker | chestnut | present | grave | 4 |
| branch | wing | cabin | engine | 1 |
| throne | surf | tower | moat | 2 |
| tank | fish | trench | medal | 2 |
| lily | newt | cat | owl | 1 |
| eel | bat | coal | ink | 1 |
| seal | coral | letter | wave | 3 |
| neon | flash | bulb | tulip | 4 |
| steam | winter | rail | station | 2 |
| queen | flower | crown | honey | 3 |
| ramp | till | basket | shelf | 1 |
| salt | pepper | bubble | lake | 2 |
| stitch | scar | wizard | school | 1 |
| desk | chalk | crayon | nail | 4 |
| window | marble | jar | milk | 4 |
| canvas | frame | cushion | sock | 2 |
| secret | treasure | coat | diary | 3 |
| cake | syrup | glue | tape | 1 |
| song | nest | chair | flock | 3 |
| tomb | kidney | sand | pyramid | 2 |
| lace | tongue | tune | heel | 3 |
| vine | cactus | leaf | shrub | 3 |
| dagger | scarf | revolver | rope | 2 |
| card | book | witch | tissue | 3 |
| rock | trophy | plate | award | 1 |
| meat | joint | ankle | roast | 3 |
| wheel | bumper | trunk | root | 4 |
| lock | nut | bolt | latch | 2 |
| inch | glow | silk | bright | 1 |
| fork | comb | rake | wrench | 4 |
| dove | snow | flour | cello | 4 |
| sword | pin | hat | scissors | 3 |
| soap | lip | bowl | stem | 1 |
| lung | actor | scalpel | surgeon | 2 |
| spoon | castle | paper | dune | 1 |
| broom | toad | word | spell | 3 |
| donkey | wolf | month | tide | 1 |
| alien | galaxy | bounty | planet | 3 |
| tire | button | clock | jacket | 4 |
| goggles | bucket | flask | scientist | 2 |
| bottle | candle | tooth | pipe | 3 |
| pole | ski | magnet | mountain | 3 |
| plain | glacier | rain | sleet | 1 |
| racket | deer | mushroom | acorn | 1 |

| **Easy semantic odd-one-out trials** | | | | |
| --- | --- | --- | --- | --- |
| ***Item 1*** | ***Item 2*** | ***Item 3*** | ***Item 4*** | ***Correct Answer*** |
| slip | fall | button | trip | 3 |
| card | present | cake | saloon | 4 |
| coffin | grave | ghost | giraffe | 4 |
| secret | yacht | boat | dinghy | 1 |
| capital | petal | stem | leaf | 1 |
| autumn | mirror | spring | winter | 2 |
| dog | chest | fish | cat | 2 |
| emerald | ruby | diamond | clock | 4 |
| violin | guitar | scientist | cello | 3 |
| apple | lemon | orange | word | 4 |
| rock | pole | paper | scissors | 2 |
| spoon | circuit | fork | knife | 2 |
| boot | slipper | revolver | shoe | 3 |
| toad | newt | chapel | frog | 3 |
| eagle | hawk | dune | owl | 3 |
| paint | canvas | brush | tractor | 4 |
| jacket | shirt | target | coat | 3 |
| dwarf | captain | sail | ship | 1 |
| needle | screen | stitch | thread | 2 |
| swing | squash | pumpkin | carrot | 1 |
| hammer | spirit | mallet | wrench | 2 |
| spade | broom | pyramid | rake | 3 |
| queen | throne | crown | drop | 4 |
| salt | veil | chilli | pepper | 2 |
| cheese | station | milk | butter | 2 |
| crayon | rat | mouse | mole | 1 |
| cotton | lace | galaxy | velvet | 3 |
| shelf | fire | candle | wax | 1 |
| sword | bow | axe | month | 4 |
| desk | cloud | rain | snow | 1 |
| lion | panther | midnight | tiger | 3 |
| bunker | trench | moat | jeans | 4 |
| soldier | surgeon | actor | kettle | 4 |
| flask | beaker | mug | car | 4 |
| eel | window | whale | shark | 2 |
| trophy | sugar | syrup | honey | 1 |
| basket | bucket | neon | box | 3 |
| cactus | saddle | bench | chair | 1 |
| domino | dice | wing | marble | 3 |
| scarf | glove | hat | grass | 4 |
| beetle | engine | wasp | cricket | 2 |
| flour | diary | letter | book | 1 |
| mansion | thorn | cabin | castle | 2 |
| frame | metre | ounce | kilo | 1 |
| scar | orchard | forest | jungle | 1 |
| tinsel | basketball | football | rugby | 1 |
| tulip | lily | nut | daffodil | 3 |
| zebra | deer | collar | donkey | 3 |
| glacier | desert | plain | towel | 4 |
| shrub | clown | flower | vine | 2 |
| jewel | coin | treasure | lake | 4 |
| dove | parrot | penguin | tower | 4 |
| font | nail | screw | pin | 1 |
| piano | star | harp | trumpet | 2 |
| turtle | seal | tissue | lobster | 3 |
| racket | spell | club | bat | 2 |
| acorn | sole | chestnut | walnut | 2 |
| wizard | magician | shell | witch | 3 |
| heel | cracker | elbow | palm | 2 |
| pillow | blanket | bell | cushion | 3 |
| wolf | fox | flash | coyote | 3 |
| soap | bubble | sink | picnic | 4 |
| pedal | banana | tomato | mushroom | 1 |
| song | psalm | tune | rail | 4 |
| panda | lizard | silk | rabbit | 3 |
| ramp | scalpel | dagger | javelin | 1 |
| heart | lung | kidney | ski | 4 |
| nest | tank | perch | flock | 2 |
| paw | till | cart | aisle | 1 |
| jar | comb | pot | bottle | 2 |
| race | lip | tooth | tongue | 1 |
| alien | plane | train | bus | 1 |
| bowl | dish | plate | craft | 4 |
| foot | mash | ankle | sock | 2 |
| branch | bark | trunk | school | 4 |
| steam | sleet | bumper | ice | 3 |
| roast | latch | fry | bake | 2 |
| tomb | tape | rope | glue | 1 |
| wheel | bounty | tire | spoke | 2 |
| award | medal | ribbon | magnet | 4 |
| joint | wine | rum | beer | 1 |
| dollar | key | handle | lock | 1 |
| arrow | shot | art | bolt | 3 |
| mountain | tide | goggles | sky | 3 |
| yard | pound | meat | inch | 3 |
| glow | bulb | bright | army | 4 |
| violet | navy | coral | root | 4 |
| surf | wave | sand | quiver | 3 |
| coal | ring | ink | chalk | 2 |
| pigeon | cow | tuna | planet | 4 |
| tax | lettuce | budget | fee | 2 |
| ivy | moss | weed | goat | 4 |
| coffee | juice | flag | soda | 3 |
| litter | triangle | square | hexagon | 1 |
| flicker | spark | flare | hoof | 4 |
| yew | oak | birch | purse | 4 |
| horse | fence | bear | pig | 2 |
| knee | beagle | hound | poodle | 1 |
| oxygen | mummy | chlorine | carbon | 2 |
| thunder | tornado | string | flood | 3 |
| rifle | bayonet | cider | grenade | 3 |
| biology | chemistry | helmet | physics | 3 |
| fifteen | million | forty | cliff | 4 |
| knuckle | pelvis | kitten | skull | 3 |
| midwife | vet | camel | doctor | 3 |
| ant | barn | fly | spider | 2 |
| marsh | peach | pond | swamp | 2 |
| jury | trial | rice | court | 3 |
| stone | poker | puzzle | chess | 1 |
| thyme | sage | guard | mint | 3 |
| jeep | roar | yell | scream | 1 |
| bridge | east | west | south | 1 |
| golf | soccer | zoo | archery | 3 |
| table | sofa | alligator | cabinet | 3 |
| shed | church | cottage | pebble | 4 |
| foal | rug | lamb | puppy | 2 |
| soprano | jail | carol | choir | 2 |
| blue | red | cave | green | 3 |
| magic | sheet | luck | charm | 2 |
| keg | angel | fairy | elf | 1 |
| vest | mule | blouse | suit | 2 |
| drill | saw | steak | pliers | 3 |
| lorry | anchor | truck | van | 2 |
| bike | noon | dawn | twilight | 1 |
| soup | jelly | pipe | cereal | 3 |
| pistol | plum | pear | grape | 1 |
| oyster | shrimp | clam | prince | 4 |
| raisin | sister | father | uncle | 1 |
| sunset | king | duchess | earl | 1 |
| pram | cannon | bullet | spear | 1 |
| hour | beef | week | year | 2 |
| silver | bird | bronze | gold | 2 |
| meadow | coach | canyon | ocean | 2 |
| dandelion | bone | rose | daisy | 2 |
| jet | tractor | ferry | ape | 4 |
| kilt | onion | cabbage | potato | 1 |
| lantern | lamp | beard | torch | 3 |
| chip | dentist | butcher | teacher | 1 |
| village | town | city | drum | 4 |
| venom | snake | bite | ball | 4 |
| moon | toe | orbit | comet | 2 |
| colony | hive | swarm | nun | 4 |
| march | soldier | troop | frost | 4 |
| valley | delta | egg | island | 3 |
| payment | debt | cost | kite | 4 |
| graph | bride | matrix | algebra | 2 |
| dart | fist | fight | punch | 1 |
| pill | duke | sultan | emperor | 1 |
| gospel | bomb | prophet | hymn | 2 |
| rainbow | steel | copper | nickel | 1 |
| gallery | statue | trout | painting | 3 |
| duvet | husband | granny | child | 1 |
| ticket | menu | leaflet | duck | 4 |
| nettle | clover | thistle | rib | 4 |
| hockey | tennis | fencing | pie | 4 |
| fog | mist | pickle | smoke | 3 |
| monsoon | storm | volcano | wig | 4 |
| feast | river | sea | creek | 1 |
| phone | radio | camera | hen | 4 |
| sweet | spice | flea | sour | 3 |

**Hard non-semantic odd-one-out items**

In each case, the correct answer is highlighted with a green border.
The first 23 trials were taken from Woolgar et al. (2010).

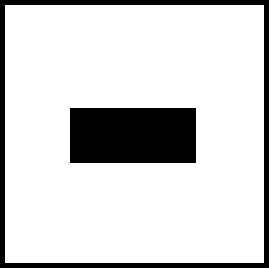

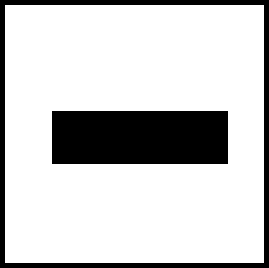

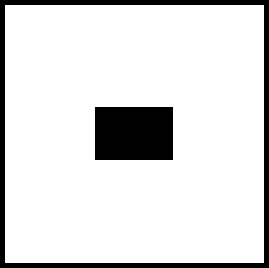

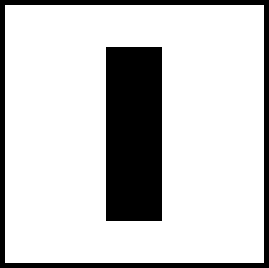

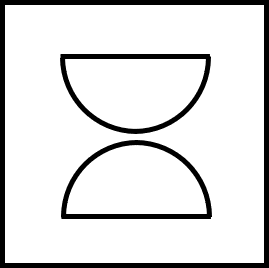

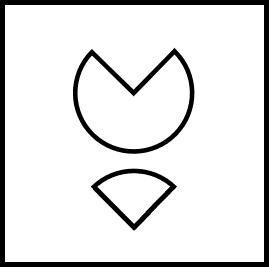

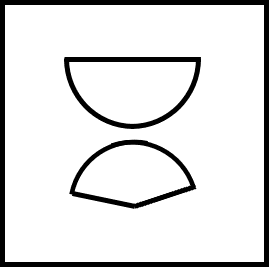

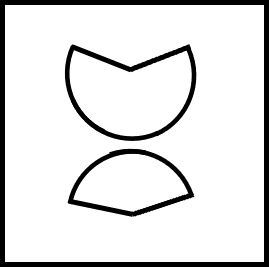

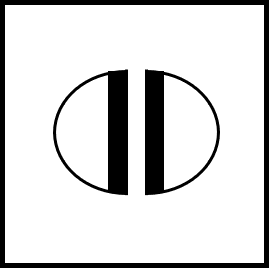

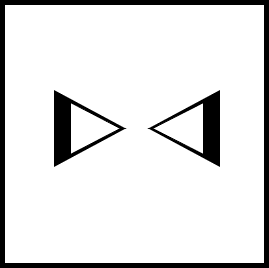

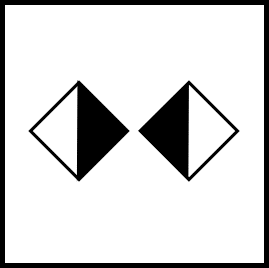

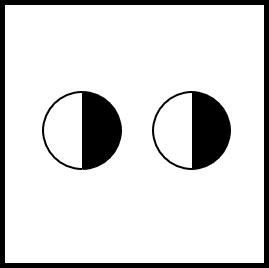

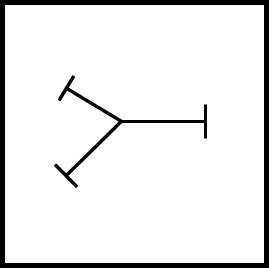

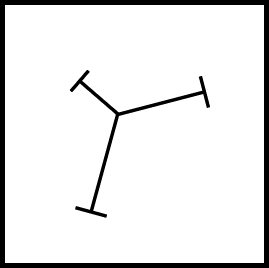

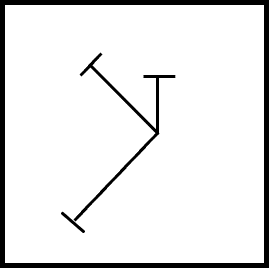

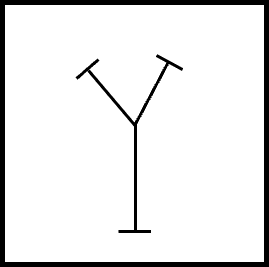

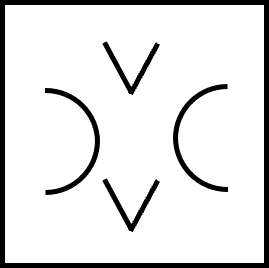

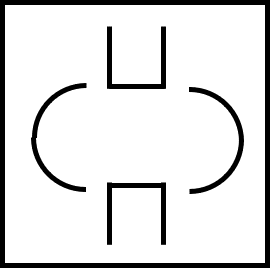

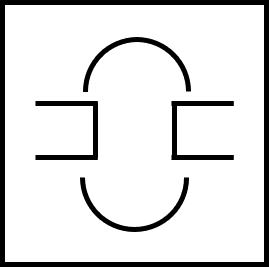

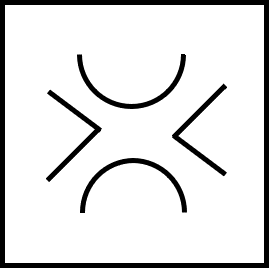

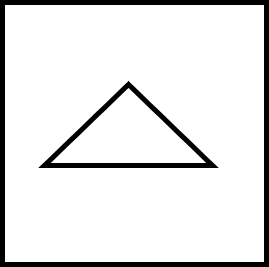

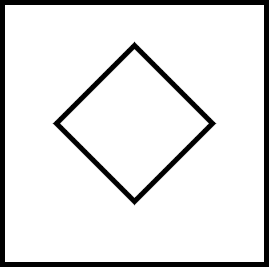

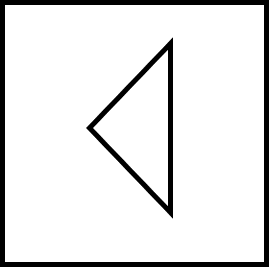

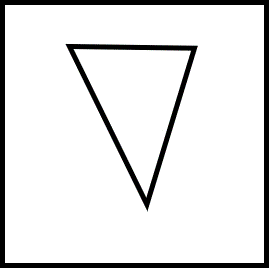

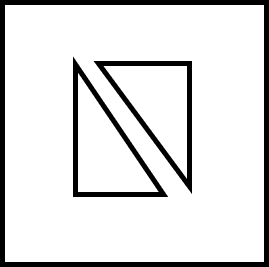

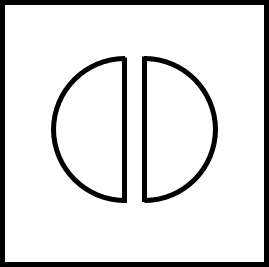

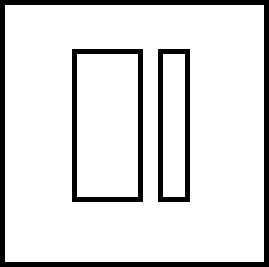

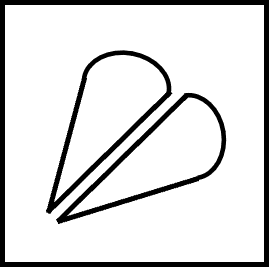

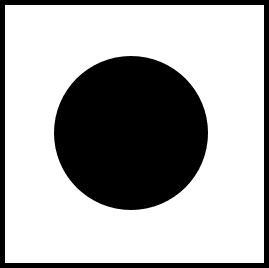

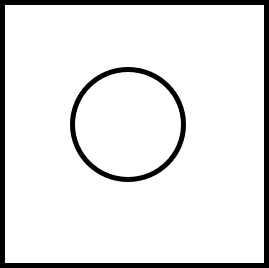

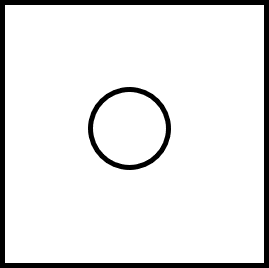

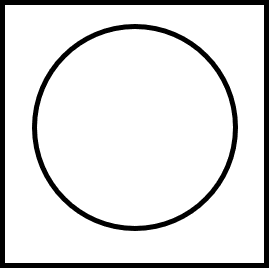

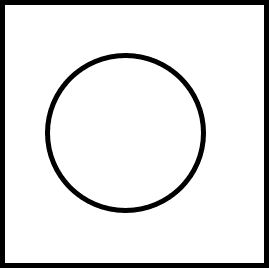

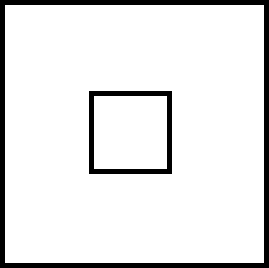

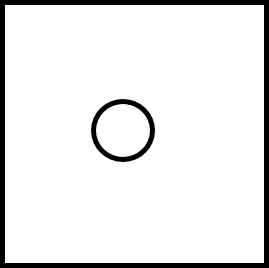

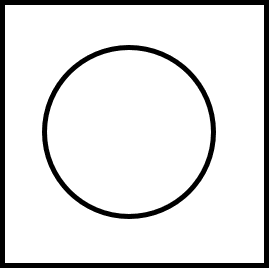

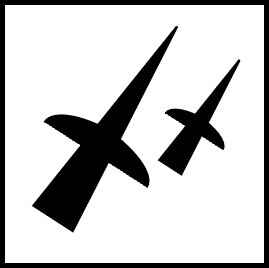

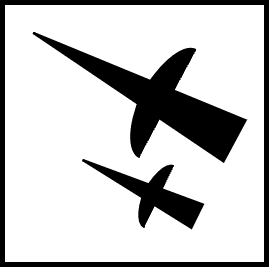

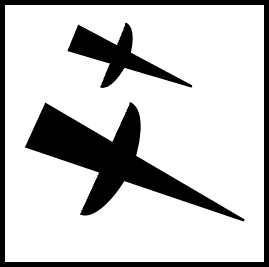

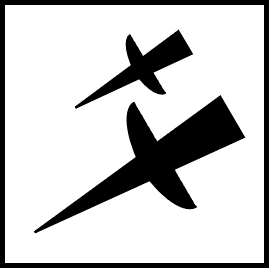

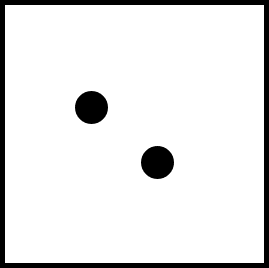

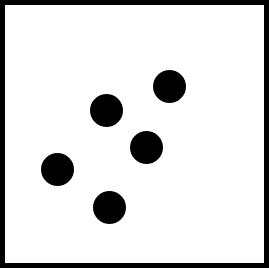

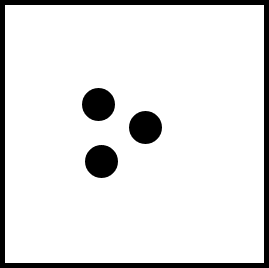

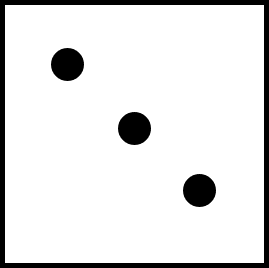

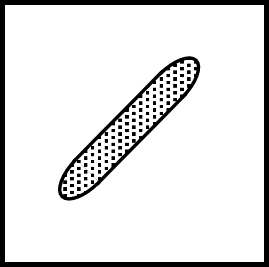

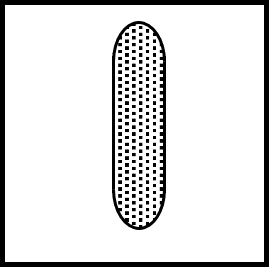

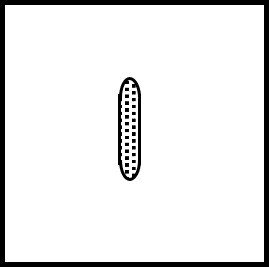

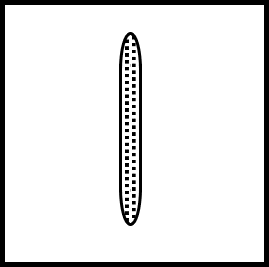

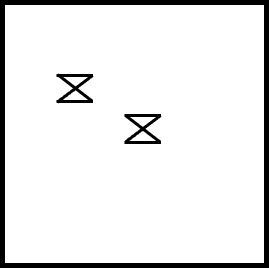

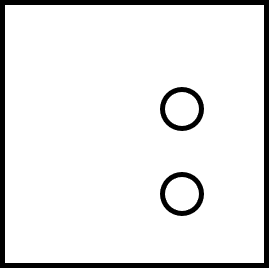

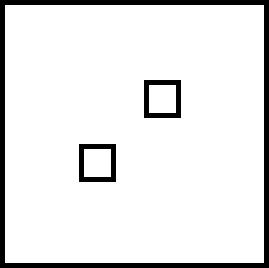

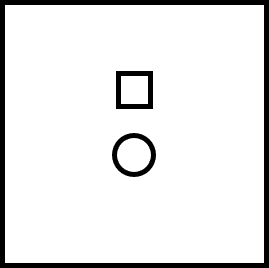

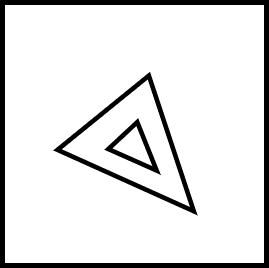

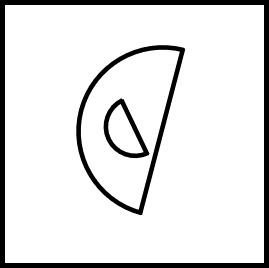

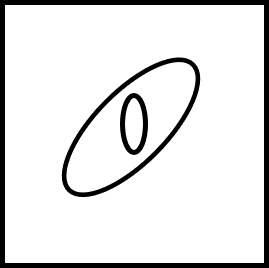

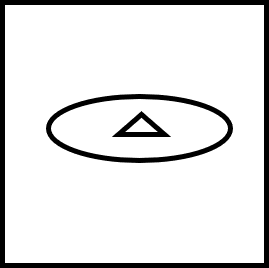

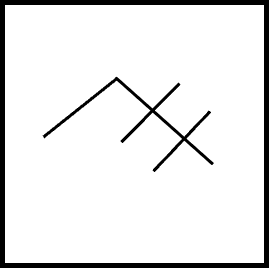

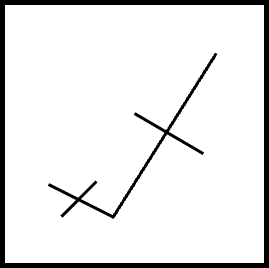

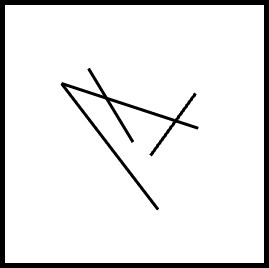

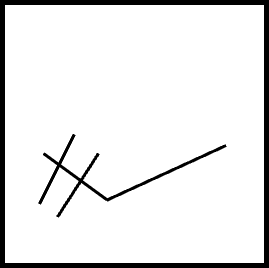

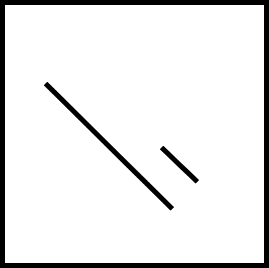

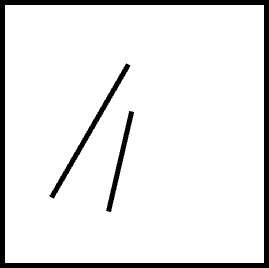

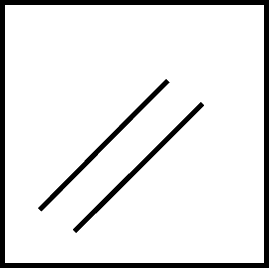

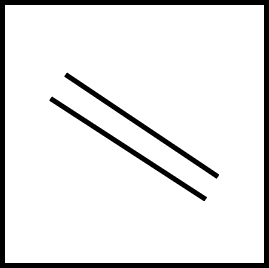

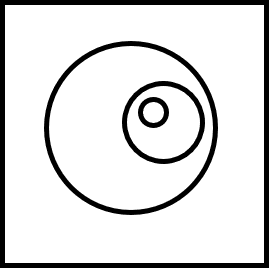

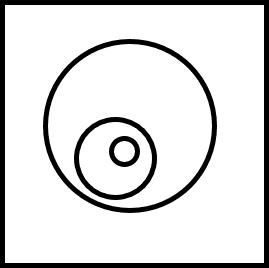

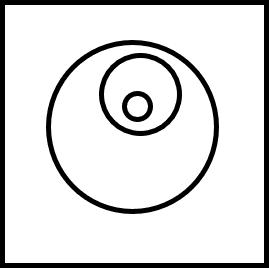

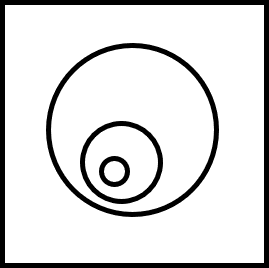

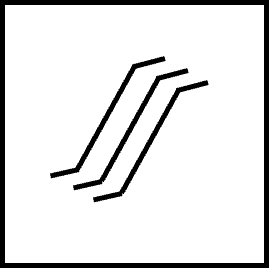

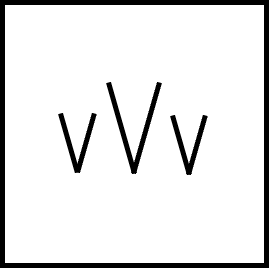

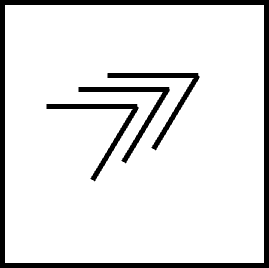

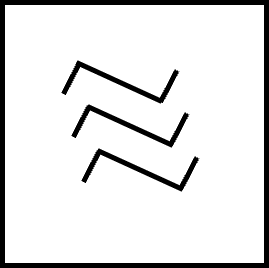

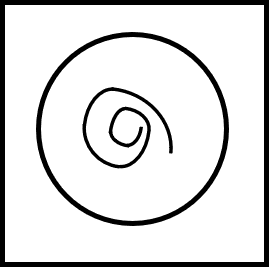

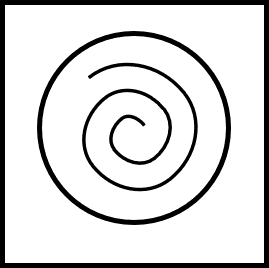

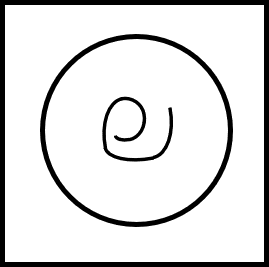

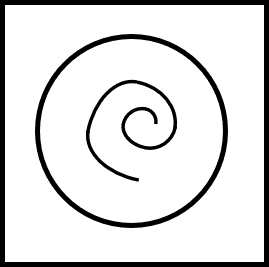

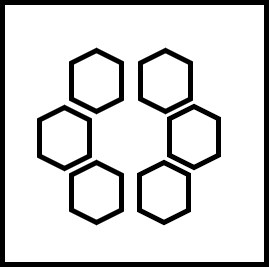

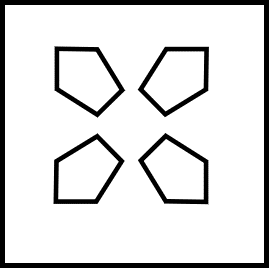

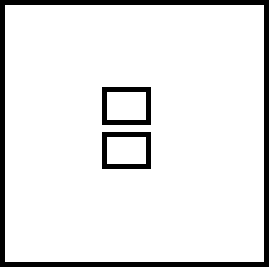

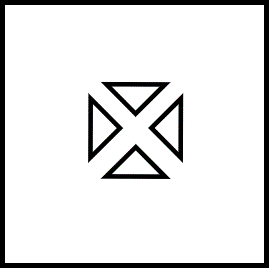

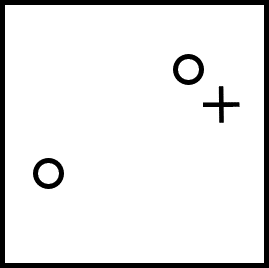

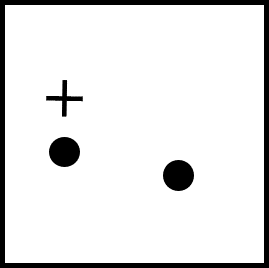

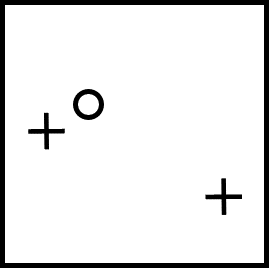

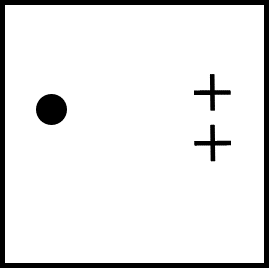

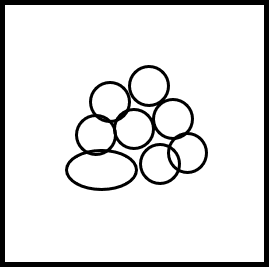

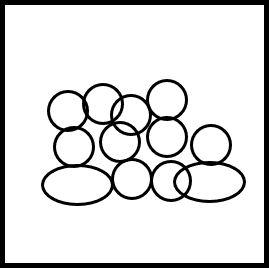

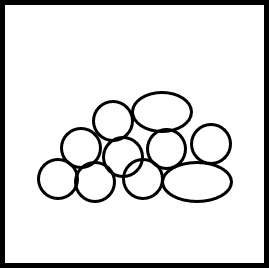

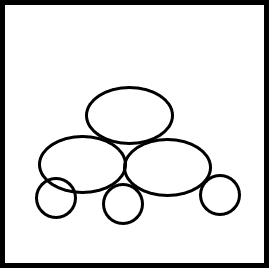

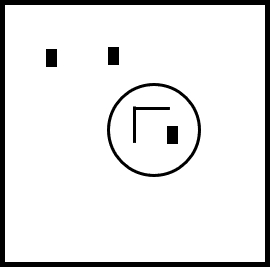

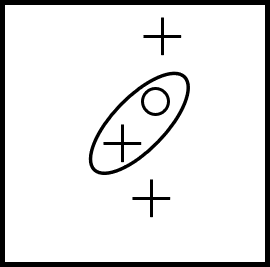

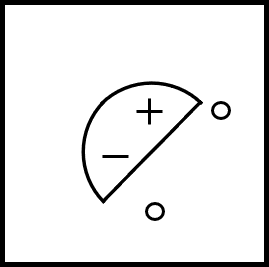

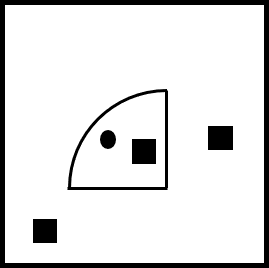

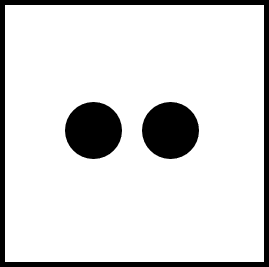

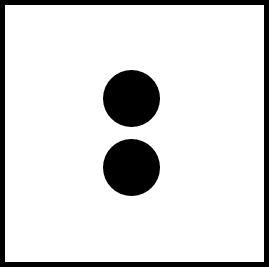

**Easy non-semantic odd-one-out items**

In each case, the correct answer is highlighted with a green border.
